# Coupling spectral and resource-use complementarity in experimental grassland and forest communities

**DOI:** 10.1101/2020.04.24.060483

**Authors:** Anna K. Schweiger, Jeannine Cavender-Bares, Shan Kothari, Philip A. Townsend, Michael D. Madritch, Jake J. Grossman, Hamed Gholizadeh, Ran Wang, John A. Gamon

## Abstract

Reflectance spectra provide integrative measures of plant phenotypes by capturing chemical, morphological, anatomical and architectural trait information. Here we investigate the linkages between plant spectral variation, spectral and resource-use complementarity that contribute to ecosystem productivity. In both a prairie grassland and a forest diversity experiment, we delineated N-dimensional hypervolumes using either wavelength-bands of reflectance spectra or foliar traits. First, we compared the hypervolume fraction unique to each species in either spectral or trait space with increasing dimensionality. Then, we investigated the association between the spectral space occupied by individual plants and their growth, as well as the spectral space occupied by plant communities and ecosystem productivity. We show that species are better distinguished in spectral space than in trait space, providing a conceptual basis for identifying plant taxa spectrally. In addition, the spectral space occupied by individuals increased with plant growth, and the spectral space occupied by plant communities increased with ecosystem productivity. Furthermore, ecosystem productivity was better explained by inter-individual spectral complementarity than by the large spectral space occupied by productive individuals. Our results indicate that spectral hypervolumes of plants can reflect ecological strategies that shape community composition and ecosystem function, and that spectral complementarity can reveal resource-use complementarity.

## 1. Introduction

Plants partition resources in space and time as a result of contrasting ecological strategies and evolutionary histories, giving rise to biochemical, structural or phenological differences that determine the optical properties of leaves and canopies. Spectral profiles of plants—defined here as the reflectance spectra of plant leaves or whole plants at high spectral resolution—are influenced by leaf traits [1-5], including pigment composition, micro- and macronutrient content, water content, specific leaf area (SLA), leaf surface properties and leaf internal structure. When measured remotely, plant spectra are also influenced by biophysical parameters of vegetation—whole-plant traits related to canopy architecture, including leaf area index, leaf angle distribution, canopy height and canopy cover [2, 6, 7]. Consequently, spectral profiles capture key differences in foliar chemistry, leaf anatomy, morphology, life history strategies and responses to environmental variation [4, 5], which have consequences for ecosystem structure and function above- and belowground [5, 8].

The optical type concept [9], published over a decade ago, posits that since neighbouring plants share resources, including light, nutrients and water, the degree of spectral complementarity in plant communities can inform us about complementary resource-use, which is, in turn, associated with ecosystem function and productivity. Complementary light use strategies have since been studied using spectral data in tropical forests [10], among evergreen and deciduous trees [11], and among prairie species [12]. Other studies have shown that dissimilarity in spectral profiles of plants correlates with their functional dissimilarity [13-17] and evolutionary divergence time [14, 15, 18, 19]; and positive relationships between measures of spectral diversity and ecosystem productivity [15] suggest a coupling between spectral differences among plants and resource-use complementarity.

Here we examine the concept of the optical types—functional differences among plants captured through optical techniques, of which spectral types represent a subset—using N-dimensional spectral hypervolumes to explore the relationship between spectral and resource-use complementarity. We define the “spectral hypervolume” as the N-dimensional space occupied by plants and delineated by spectral axes—spectral bands or other expressions of spectral variation—along which plants can vary.

Plant spectral hypervolumes can be quantified across biological scales from individual plants to species, lineages or even plant communities. The spectral hypervolume occupied by an individual plant can be quantified by leaf spectra from that plant. Its size represents intra-individual spectral variation [20] and is conjectured to be indicative of the range of environmental conditions experienced by the plant, including gradients of light, wind and temperature, as well as variation in pathogen and herbivore pressure within the canopy. Greater variation in environment and in leaf spectra within a plant give rise to a larger spectral hypervolume, which is in turn, we hypothesize, positively correlated with plant growth and biomass. The spectral hypervolume occupied by a species can be quantified by spectral profiles of individuals from that species within a certain ecosystem or distributed across ecosystems. The size of species’ spectral hypervolumes represents intraspecific spectral variation; distances among the spectral hypervolumes of different species are correlated with species’ spectral dissimilarities; and the degree of overlap among species’ spectral hypervolumes is indicative of their shared spectral characteristics, which are associated with shared functional attributes [2, 9, 15] and shared ancestry [5, 14, 15, 19]. The spectral hypervolume occupied by a plant community can be quantified by the spectral profiles of individuals and species within that community. The size of the spectral hypervolume occupied by a plant community represents the variation among spectral types within that community, or its spectral diversity, which in turn is hypothesized to be indicative of the community’s taxonomic, functional and phylogenetic diversity [15, 21]. Greater variation among plant spectra within a community gives rise to a larger community hypervolume. Overlaps among the spectral hypervolumes of plant communities indicate similarity in resource-use patterns among these communities, while distances among spectral hypervolumes of plant communities are hypothesized to predict the degree of dissimilarity in resource-use patterns among them and the dissimilarity in environmental conditions they experience.

We define “spectral complementarity” as the separation of spectral hypervolumes of plants within multidimensional spectral space. Spectrally complementary species, which represent contrasting spectral types, can be assumed to use resources differently, including light, nutrients and water, because resource-use strategies of plants are reflected by their foliar chemistry, anatomy and morphology [22, 23], all of which influence the spectral response following the physics of light absorption and scattering [2]. From an individual perspective, spectral complementarity means that leaves from different parts of the canopy complement each other in terms of resource-use, for example, through foliar adaptations in response to light gradients within canopies [20, 24]. From a stand or community perspective, spectral complementarity means that different individuals or species partition resources which allows them to compete less with each other and use the total resource pool together more completely [25-27] leading to a positive relationship between spectral complementarity, total resource-use and productivity.

The framework of the N-dimensional hypervolume has a long history in ecology. Proposed by Hutchinson in connection with the N-dimensional niche concept [28], hypervolumes have been used frequently to quantify the multivariate environmental or trait space occupied by organisms, including plants [29], to describe the niches or multivariate functions of species or broader clades. They have also been used to describe plant communities or biomes [30, 31]. Notably, in remote sensing there are well-established analogous concepts, including spectral endmembers, that are related to the positioning of reflectance spectra in multi-dimensional spectral space which are frequently used for feature extraction and classification [32]. The basic notion is that to differentiate and classify objects from remotely sensed imagery into groups, these groups need to show sufficient separation in spectral space for distinct clusters to emerge.

Here, we use spectral hypervolumes to assess variation among plants at different biological scales—individuals, species and communities—to understand the consequences of these dissimilarities for ecosystem function. Our overarching hypothesis is that the spectral complementarity of plants is indicative of their ecological complementarity in terms of resource-use. Based on this main hypothesis we test three specific hypotheses using data collected either in a prairie grassland diversity experiment— Biodiversity II (BioDIV) [33]—or a tree diversity experiment—Forest and Biodiversity I (FAB) [34]—or both. First, we expect the high dimensionality of spectral data to detect plant species differences more readily than commonly measured foliar traits, many of which may be similar among species growing in a common abiotic environment. Consequently, we expect spectral hypervolumes to overlap less among species than trait hypervolumes (BioDIV). If so, differentiation of species in spectral space has the potential to indicate spectral complementarity and provides a conceptual basis for spectral identification of plants [18, 35, 36] grounded in ecological theory. Second, we hypothesize that greater spectral variation among leaves within focal plants, which results in larger individual spectral hypervolumes, will be associated with greater individual growth (FAB). Finally, we hypothesize that plant communities that occupy greater total spectral space—either due to spectral complementarity among species or due to individuals that occupy large spectral hypervolumes or both—will be associated with more productive ecosystems (FAB and BioDIV).

## 2 Methods

### (a) Study sites

Both BioDIV [33] and FAB [34], which is part of IDENT and TreeDivNet, are located at the Cedar Creek Ecosystem Science Reserve (East Bethel, MN). We used leaf spectra and foliar traits of 902 individuals from 14 grassland–savannah perennials sampled in 35 plots in BioDIV and aboveground biomass determined in the same plots in July 2015; and leaf spectra, plant height and diameter measurements of 537 individuals from 12 tree species sampled in 68 plots in FAB in July 2016. The species sampled in BioDIV were: *Achillea millefolium* L. (49 ind.), *Amorpha canescens* Pursh (28 ind.), *Andropogon gerardii* Vitman (162 ind.), *Asclepias tuberosa* L. (70 ind.), *Lespedeza capitata* Michx. (99 ind.), *Liatris aspera* Michx. (49 ind.), *Lupinus perennis* L. (121 ind.), *Panicum virgatum* L. (49 ind.), *Petalostemum candidum* (Willd.) Michx. (28 ind.), *Petalostemum purpureum* (Vent.) Rydb. (52 ind.), *Petalostemum villosum* Nutt. (42 ind.), *Schizachyrium scoparium* (Michx.) Nash (76 ind.), *Solidago rigida* L. (50 ind.) and *Sorghastrum nutans* (L.) Nash (27 ind.). The species sampled in FAB were: *Acer negundo* L. (30 ind.), *Acer rubrum* L. (47 ind.), *Betula papyrifera* Marshall (44 ind.), *Juniperus virginiana* L. (39 ind.), *Pinus banksiana* Lamb. (47 ind.), *Pinus resinosa* Aiton (52 ind.), *Pinus strobus* L. (47 ind.), *Quercus alba* L. (42 ind.), *Quercus ellipsoidalis* E.J.Hill (49 ind.), *Quercus macrocarpa* Michx. (50 ind.), *Quercus rubra* L. (39 ind.) and *Tilia americana* L. (51 ind.).

### (b) Spectral data

We measured leaf spectra using two portable field spectrometers (SVC HR-1024i, Spectra Vista Corp., Poughkeepsie, NY; and PSR+ 3500, Spectral Evolution Inc., Lawrence, MA) and the associated leaf-clip assemblies (LC-RP PRO, Spectra Vista Corp.; and PSR+ 3500 leaf clip, Spectral Evolution Inc.) covering the wavelength range from 350 nm to 2500 nm in 1024 spectral bands. We used the SVC and PSR+ instruments for spectral measurement of herbaceous and tree species, respectively. To characterise one individual spectrally, we measured the reflectance of either three or five mature, healthy leaves per individual depending on plant height. Spectra were automatically corrected for dark current and stray light, and referenced against the Spectralon^®^ 99% reflectance standard disc (Labsphere, North Sutton, NH) of the leaf clip approximately every 10 minutes. Spectral data processing included correcting discontinuities at the sensor overlap regions between the Si and first InGaAs sensor (around 1000 nm) and between the first and second InGaAs sensor (around 1900 nm), removing noisy regions at the beginning and end of the spectrum, and interpolating spectra to 1 nm resolution. For spectral processing we used the spectrolab [37] package in R [38].

### (c) Foliar traits and measures of individual plant growth

For all individuals of grassland–savannah perennials measured with the leaf clip, we estimated the following foliar traits using partial least squares regression (PLSR) models [39] previously developed from chemical assays of leaf tissue samples and corresponding leaf spectra [15]: leaf nitrogen, carbon, non-structural carbohydrates, hemicellulose, cellulose and lignin concentration (%), and the content of chlorophyll a and b, beta-carotene, lutein, violaxanthin, antheraxanthin and zeaxanthin pigments. We summarised chlorophyll content as chlorophyll a plus chlorophyll b, expressed as μmol m^-2^; and we expressed beta-carotene and lutein content, and the size of the xanthophyll pigment pool (VAZ = violaxanthin plus antheraxanthin plus zeaxanthin) as ratios relative to chlorophyll content to indicate contrasting photosynthetic behaviour and photoprotective capacity among plants [16]. The chemical methods used were in brief: combustion–reduction elemental analysis (TruSpec CN Analyzer; LECO, St. Joseph, MI) to determine leaf nitrogen and carbon concentrations; sequential digestion (Fiber Analyzer 200; ANKOM Technology, Macedon, NY) to determine carbon fractions; and high-performance liquid chromatography (HPLC) to determine pigment contents (for details see [15]). We tested the degree to which species differed regarding their foliar traits using Tukey’s honest significant difference (HSD) post-hoc tests and the R package agricolae [40]. Further, since we expected species separability to increase with phylogenetic distance [15], we tested for phylogenetic signal of each trait using Blomberg’s K statistic [41] as implemented in the R package picante [42] and the phylogeny reconstructed by Kothari et al. (2018) [12] with one missing species (*Petalostemum candidum* (Willd.) Michx.) added manually using phytools [43]; for details see electronic supplementary material, table S1. In the FAB experiment, we used the biophysical parameters tree height (cm) and basal diameter (mm) as measures of individual growth (for details see [34]).

### (d) Species hypervolumes in spectral and trait space

We quantified spectral hypervolumes using kernel density estimation, which can describe complex shapes in high dimensionalities, measure the volume of these shapes and perform set operations (union, intersection, difference) on multiple shapes [30, 31]. We calculated the hypervolume fraction unique to each of the 14 BioDIV species in spectral space and trait space of increasing dimensionality using the R package hypervolume [32]. We randomly selected between 2 and 21 spectral bands and between 2 and 10 foliar traits as axes delineating species’ spectral and trait hypervolumes, and repeated selections 50 times each; leaf traits were centred and scaled. We projected all individuals into the resulting spectral and trait spaces, and calculated the fraction of the hypervolume unique to each species (i.e., the hypervolume that is occupied by the focal species and not overlapped by any other species). Since it is difficult to show more than three dimensions in one graph, we used linear discriminant analysis (LDA) to illustrate species dissimilarity in n-dimensional spaces delineated by the main axes of spectral and trait variation. In our case, linear discriminants (LDs) are linear combinations of all band-wise reflectance and trait values, respectively, which re-project observations into a new coordinate system while maximising the differences between groups; our grouping variable was species identity. For LDA we used the R package MASS [44]; for interactive 3D graphics illustrating species hypervolume shifts with changes in community diversity we used plotly [45].

### (e) Species identification models

Since we are also interested in whether the degree of species differentiation in spectral and trait space is tied to the accuracy of species classification models, we tested the degree to which plant species in BioDIV can be correctly identified based on their spectra and foliar traits with partial least squares discriminant analysis (PLSDA) as implemented in the R package caret [46]. We used random draws of 20 individuals per species for model training and the remaining data for validation and for evaluating model fit. We performed 100 PLSDA model iterations using new random draws of training samples and selected the optimal number of components based on the minimum of the root mean squared error of prediction (RMSEP) for the validation samples. We tested for significant differences in RMSEP values among the number of components using Tukey’s HSD test and used the smaller number of components when models performed similarly (p > 0.05). We investigated which wavelengths and foliar traits contributed most to species separability using PLSDA loadings to determine if spectral differences among species represent differences in traits representing physiological function or resource-use strategies. All statistics and graphs are based on the validation results.

### (f) Spectral space occupied by individuals and plant communities

We tested the degree to which the spectral space occupied by individual plants predicts plant growth by fitting regression models between the spectral space occupied by individual trees sampled in FAB and tree height (cm) or basal diameter (mm), which served as proxies for biomass. Trees in FAB were planted a similar time and size such that plant size provided a measure of plant biomass closely related to growth rate and productivity. We did not perform the same test in BioDIV because no measures of individual plant growth are available in this experiment.

Next, we tested the degree to which the spectral space occupied by plant communities predicts aboveground ecosystem productivity. In FAB, we used overyielding, the excess biomass produced by mixed species plots compared to what would be expected based on monoculture yields, as a measure of the net biodiversity effect (NBE). In addition, we partitioned the NBE into complementarity (CE) and selection effects (SE), following Loreau and Hector (2001) [27], to test the degree to which spectral complementarity among individuals and individuals occupying large spectral spaces contribute to overyielding. Biomass for NBE, SE and CE calculations was determined from allometrically derived incremental stem biomass (kg y^-1^; for details see [34]). In BioDIV, we used biomass (g m^-2^, dry weight) determined in clipstrips as a measure of aboveground net primary productivity. We did not calculate and partition the NBE in BioDIV because monocultures are not replicated in this experiment and calculations of the NBE and its components depend on estimates of mean monoculture yields. Since hypervolume size is known to be positively correlated with sample size [29], we used 12 randomly selected individuals per plot to calculate the spectral space occupied by plant communities in BioDIV, resulting in a total of 30 communities used for analysis. In FAB, we used nine randomly selected individuals per plot, resulting in a total of 68 communities used for analysis. We reduced data dimensionality to the first three principal component (PC) axes, which explained more than 98% of the total spectral variation, calculated the spectral space occupied per community using the R package hypervolume [31] and tested the association between the spectral space occupied by plant communities and community productivity using linear regression models. In addition, we tested whether the spectral space occupied by plant communities increases with species richness (see electronic supplementary material, appendix S1).

## 3 Results

### (a) The degree of species differentiation in spectral and trait space

Species’ spectral hypervolumes were more distinct than their trait-based hypervolumes calculated from the 10 chemical and structural foliar traits measured in our study. The fraction of the hypervolume space unique to each of our 14 species of grassland–savannah perennials increased with the dimensionality of spectral and trait space (figure 1). However, while each focal species occupied a hypervolume that was at least 90% unique to the species after including 15 randomly selected spectral bands as axes, not all focal species reached the same level of uniqueness in trait space after including all 10 foliar traits as axes (figure 1; electronic supplementary material, tables S2 and S3). We confirmed that species’ spectral hypervolumes were more distinct than their trait hypervolumes by projecting species’ positions along the main axes of variation in spectral or trait space. In spectral space, all non-graminoid species clearly separated along the first four linear discriminant axes (LDs), and LDs 11 and 12 separated the graminoids. In trait space, however, only few species formed distinct clusters and we found no combination of LDs that separated the four graminoids species from each other (electronic supplementary material, figure S1). These overlaps are likely due to the substantial degree of intraspecific variation across all measured traits and the lack of a particular trait or traits that differed significantly among all species (electronic supplementary material, figure S2, table S4). This is confirmed by the high degree of overlap among species hypervolumes in two-dimensional trait space (electronic supplementary material, figure S3 right); although it is worth noting that broader taxonomic groups clustered away from other species along certain trait axes, including legumes along carbon content, and graminoids along nitrogen content and carbon fraction axes. Species hypervolume overlap was also pronounced in two-dimensional spectral space delineated by the 10 most variable spectral bands, but spectrally distinct species already started emerging (electronic supplementary material, figure S3 left).

**Figure 1.**
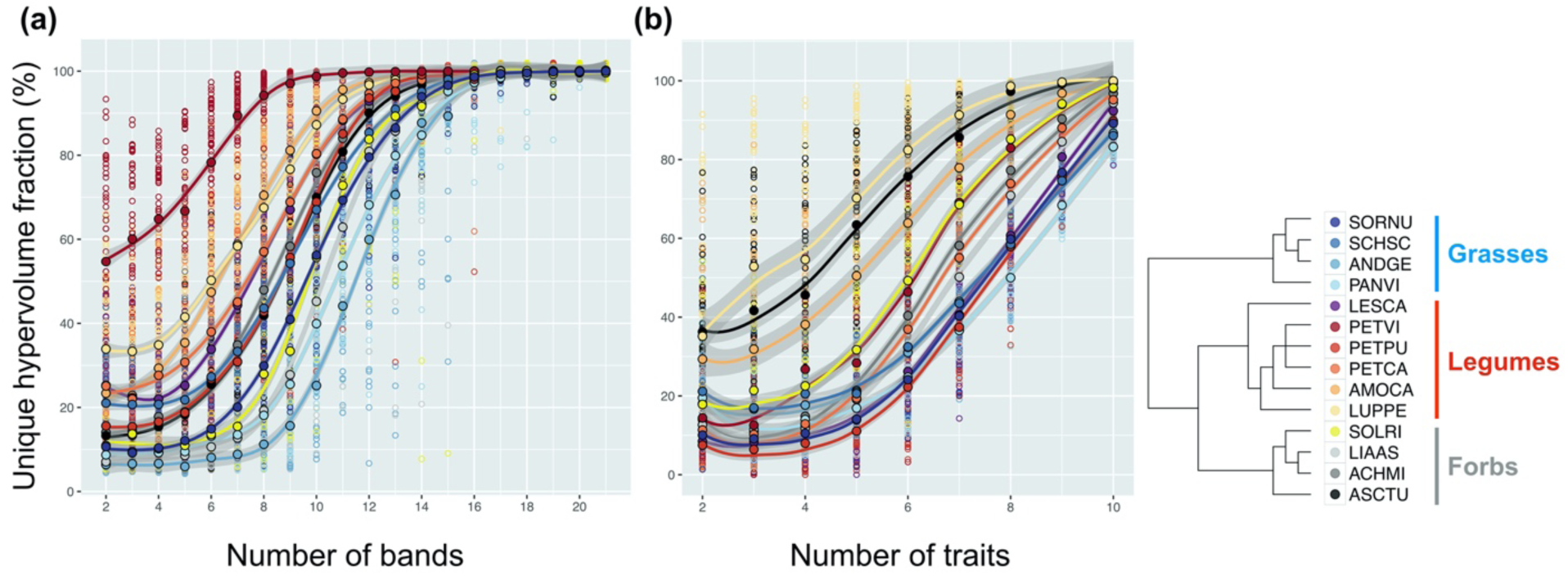
The unique hypervolume fraction per species increases with increasing dimensionality of *(a)* spectral space and *(b)* trait space. Curves represent second order polynomials fitted to 50 estimates of species hypervolume sizes calculated with increasing number of dimensions (2-21 spectral bands and 2-10 foliar traits, respectively); 95% confidence interval are shown in grey. The species are *Sorghastrum nutans* (L.) Nash (27 ind., SORNU), *Schizachyrium scoparium* (Michx.) Nash (76 ind., SCHSC), *Andropogon gerardii* Vitman (162 ind., ANDGE), *Panicum virgatum* L. (49 ind., PANVI), *Lespedeza capitata* Michx. (99 ind., LESCA), *Petalostemum villosum* Nutt. (42 ind., PETVI), *Petalostemum purpureum* (Vent.) Rydb. (52 ind., PETPU), *Petalostemum candidum* (Willd.) Michx. (28 ind., PETCA), *Amorpha canescens* Pursh (28 ind., AMOCA), *Lupinus perennis* L. (121 ind., LUPPE), *Solidago rigida* L. (50 ind., SOLRI), *Liatris aspera* Michx. (49 ind., LIAAS), *Achillea millefolium* L. (49 ind., ACHMI), *Asclepias tuberosa* L. (70 ind., ASCTU). Phylogenetic relationships among species are displayed on the right; graminoids are coded in blue, legumes in purple-red-orange and forbs in yellow-grey-black colours; for statistics see electronic supplementary material, tables S2 and S3.

### (b) Species identification models

Species identification models based on spectra (65% - 98% accuracy per species) outperformed species identification models based on traits (47% - 97% accuracy per species; figure 2; electronic supplementary material, figure S4, table S5). This was consistent with the smaller overlap (figure 1) and greater distance (electronic supplementary material, figure S1) between species’ hypervolumes in spectral space compared to trait space. The spectral bands contributing most to species’ separability aligned with absorption features related to leaf chlorophyll, carotenoid, lignin and protein content [3] (electronic supplementary material, figure S5*a*). These foliar traits also contributed most to species separability in trait space (electronic supplementary material, figure S5*b*) and all of them, except for chlorophyll content, showed evidence of phylogenetic signal (electronic supplementary material, table S2, figure S6) highlighting the potential importance of these traits for species identification across ecosystems.

**Figure 2.**
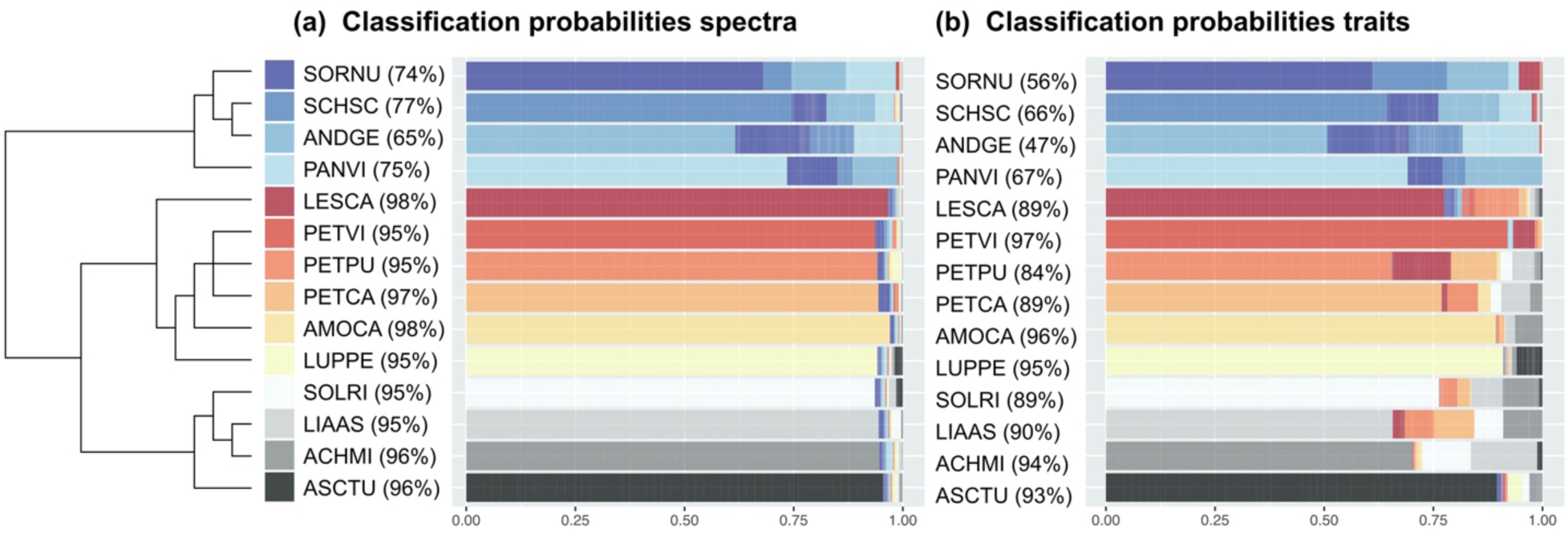
Probabilities that each species is classified correctly or incorrectly classified as another species based on partial least squares discriminant analysis (PLSDA) models using *(a)* spectra and *(b)* foliar traits of species sampled in the BioDIV experiment. The percentages of correctly classified samples per species are shown in parentheses. PLSDA also assigns to every sample the classification probabilities to belong to each class (i.e., species). Coloured bars that match the species colour, shown at the tip of the phylogeny, indicate the averaged probability that the species was correctly classified. Bars that show a different colour indicate the averaged probability that the species was misclassified as a different species. The graph shows model validation results, i.e., n = the number of individuals per species minus 20 individuals for model calibration; for species abbreviations and number of individuals per species see figure 1.

### (c) Individual hypervolume size is associated with plant biomass

The spectral space occupied by individual trees—a measure of intra-individual spectral variation—was positively correlated with tree height and diameter (figure 3), both of which predict biomass and are closely correlated with productivity—the increase in biomass per unit time. This is likely because trees that grow more tend to have larger, more complex canopies and higher foliar plasticity than trees that grow less. Over time, taller trees seem to be able to sustain and perhaps even increase the benefits gained from harnessing a more diverse light environment (electronic supplementary material, figure S7), pointing towards size-asymmetric (size-dependent) light competition [47] among the trees in the FAB experiment. The spectral space occupied by individuals was more closely associated with tree diameter than tree height. One explanation is that once trees are taller than their neighbours, it may be more advantageous to invest in mechanical stability and horizontal canopy extension than in vertical growth to maximise light interception.

**Figure 3.**
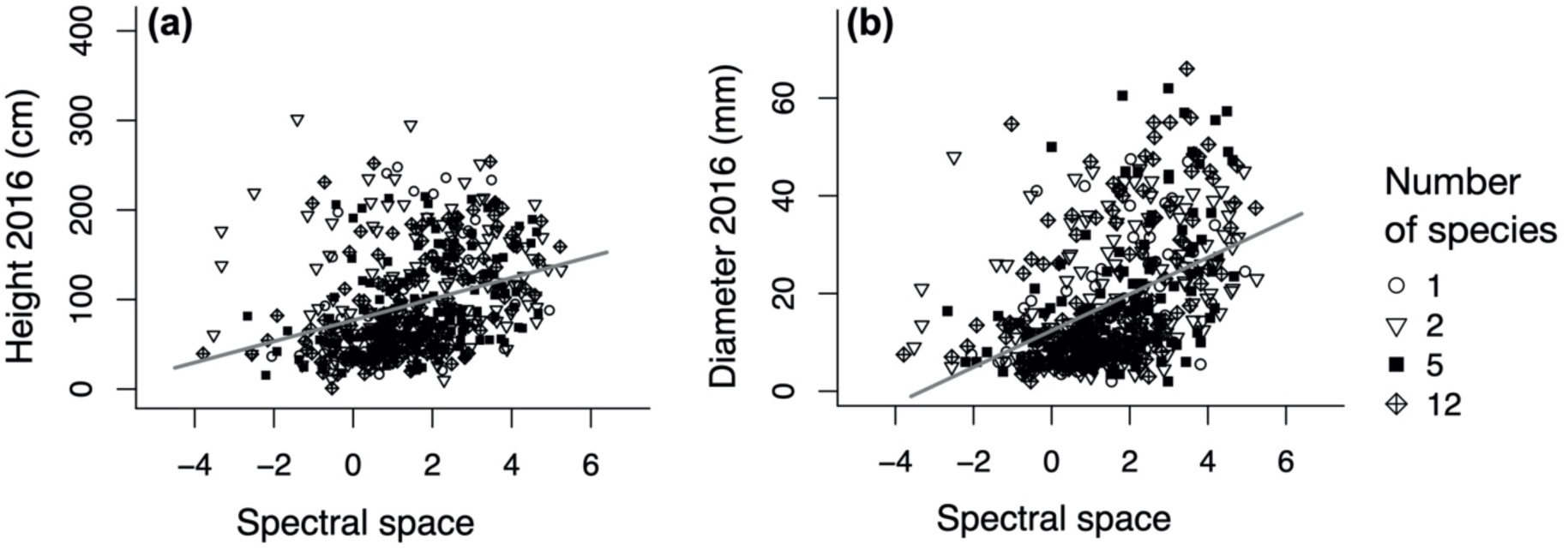
The spectral space occupied by individual trees in the FAB experiment predicting *(a)* tree height (n = 532, r^2^ = 0.11, F_1,530_= 63.2, P < 0.001) and *(b)* stem diameter (n = 532, r^2^ = 0.20, F_1,530_ = 135.5, P < 0.001) in 2016. Spectral hypervolume sizes are log-transformed; the number of species per plot is indicated with different symbols.

### (d) More productive communities occupy larger spectral hypervolumes than less productive communities

The spectral space occupied by communities in the BioDIV experiment explained 44% of the total variation in aboveground productivity (figure 4). Likewise, the spectral space occupied increased with the number of species per plant community (electronic supplementary material, figure S8*a*). In FAB, we used overyielding as a measure of the net biodiversity effect (NBE) [27]; and we partitioned the NBE into its two components, complementarity (CE, the positive effects of diverse resource-use strategies and positive interactions among plants on productivity) and selection effects (SE, the effect of the dominance of productive species in mixtures on productivity). The spectral space occupied by tree communities in FAB explained 37% of overyielding (figure 4*b*) and increased with the number of species per community (electronic supplementary material, figure S8*b*). Partitioning the NBE into its two components revealed a positive relationship between the spectral space occupied by communities and complementarity, while the association with the selection effect was negative (electronic supplementary material, figure S9). In other words, compared to less productive communities more productive communities did not, on average, harbour more highly productive individuals which occupy larger spectral spaces, but rather more spectrally complementary species that collectively contributed to the large spectral space occupied by these communities.

**Figure 4.**
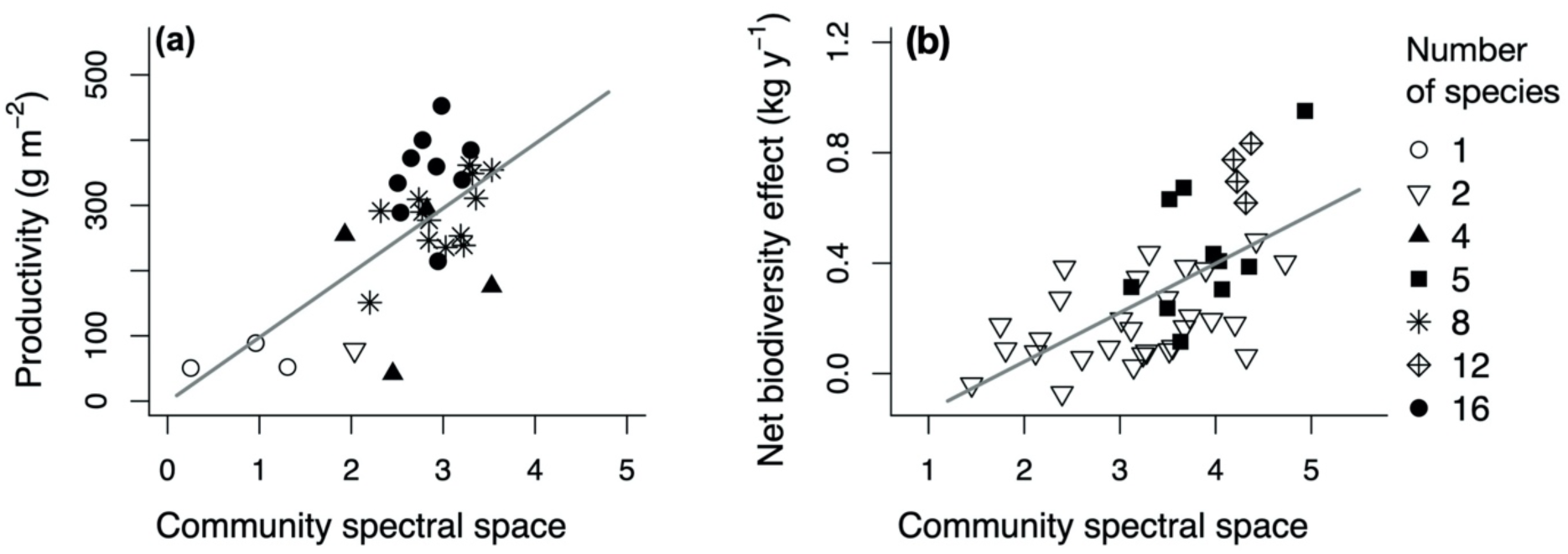
Spectral hypervolume occupied by plant communities predicting *(a)* aboveground productivity in the BioDIV experiment (n = 30, r^2^ = 0.44, F_1,28_ = 22.6, P < 0.001) and the *(b)* net biodiversity effect (NBE) in the FAB experiment (n = 43, r^2^ = 0.37, F_1,41_= 24.3, P < 0.001). The number of species planted per plot is indicated with different symbols; spectral hypervolume sizes are log-transformed.

## 4 Discussion

Plant spectra provide integrative measures of plant phenotypes. Quantification of plant N-dimensional spectral hypervolumes offers a novel and effective way to assess variation among plant phenotypes associated with resource-use complementarity that is predictive of plant growth and ecosystem productivity. We find that that greater spectral complementarity—the degree to which individuals and species occupy distinct hypervolumes in spectral space—is associated with greater resource capture and growth in focal plants, and greater ecosystem productivity in communities with larger spectral hypervolumes. Consequently, we posit that spectral complementarity provides a measure of resource partitioning in plant communities. The multidimensional separation of plants in spectral space also provides a conceptual basis for distinguishing plant taxa spectrally. Many functional traits of plants, including traits specific to organs that interact minimally with light, such as roots and seeds, may not influence the spectral signal directly. However, plant spectra integrate many aspects of plant form and function, including biochemical, anatomical and morphological traits [5, 9], which are sometimes expressed in coordination across the whole plant [48]. For example, as plants use light to power photosynthesis, their physiology responds in a coordinated way to solar radiation and atmospheric conditions (e.g., vapour pressure deficit), and also to belowground resources (e.g., water and nutrients). This leads to responses throughout the canopy including short- and mid-term changes in pigment pool sizes [16], hydraulic properties [49] and adaptations in growth form, providing a strong basis for using spectra as measures of plant function.

### (a) The degree of species’ differentiation in spectral space

Projecting species into trait and spectral spaces with increasing dimensionality gradually reduced the overlap among species (figure 1). However, species trait hypervolumes contracted to a lesser degree than their spectral hypervolumes, such that models identifying species using spectra outperformed those using foliar traits (electronic supplementary material, table S5). The degree of species differentiation in spectral space is not surprising given the high dimensionality of spectral data. In addition, spectral measurements may provide a more objective metric of traits or trait proxies, as they, when performed properly and consistently, lack some of the errors of extractive measurements. As integrative measures of plant phenotypes, spectra of plants are expected to describe the total dissimilarity among plants resulting from their differences in biochemistry, anatomy and morphology more completely than a number of selected traits [5, 15, 50]. To some extent, this effect could be due to redundancy in our trait measures. In our case, light gradients are probably the dominant source of environmental variation, and all leaf traits measured in our study are to some degree influenced by variation in light. For instance, the ratio of chlorophyll and carotenoid pigment levels reflects biochemical acclimation to stress under different light environments [12, 51]. Likewise, the contents of different carbon fractions are tied to morphological adaptations, such as leaf thickness and SLA, to light gradients within canopies [24]. In this way, what we think of as multiple traits can also be thought of as different proxies for the same or overlapping traits [7].

Misclassifications occurred more often among closely related than among distantly related species (figure 2; electronic supplementary material, figure S4*a*), likely due to the similarity in foliar traits (electronic supplementary material, tables S1 and S4, figure S6) and spectra among close relatives [15, 18]. Investigating the spectral regions that contribute most to species separability confirmed a tight coupling between spectral and ecological complementarity in terms of resource-use. The spectral regions that contributed most to species separability aligned with absorption features for proteins and lignin content [3] (electronic supplementary material, figure S5*a*), leaf traits associated with the trade-off between fast and slow return on investment [22], highlighting the relationship between differences in resource-use strategies and spectral complementarity. In addition, chlorophyll content and xanthophyll cycle pigment pool size contributed to species separability, suggesting that prairie ecosystems harbour species with different strategies for light capture and photoprotection [12], which has also been found in tropical forests [10]—both are examples of linkages between spectral complementarity and resource-use. These sets of traits also relate to canopy structure, so they may be associated with architectural differentiation that influences complementarity in spectra and plant traits at the canopy scale [52].

### (b) Spectral hypervolumes occupied by individuals and plant communities

The positive relationships between intra-individual spectral variation, and tree height and diameter (figure 3) confirm our hypothesis that plants occupying more spectral space have greater growth. Higher growth rates create stronger light gradients within canopies driving plasticity in foliar traits, including SLA, pigments and nitrogen content [53, 54], which all influence the spectral response. Foliar plasticity allows trees to minimise the costs of light capture in internal leaves relative to the benefits, which makes light capture more efficient overall than if there were no plasticity under the same conditions. However, while tree growth in FAB was positively correlated with intra-individual spectral variation, the size of the spectral hypervolumes occupied by productive individuals did not explain the association between community productivity and the size of the spectral hypervolume occupied by communities (figure 4; electronic supplementary material, figure S9). In our case, it appears that the size of the spectral hypervolume occupied by productive plant communities is largely attained by spectral complementarity, which contributes to contrasting spectral patterns among evergreen and deciduous boreal trees [11], in prairie species [12] and in dry tropical forests [16]. We posit that the large spectral space occupied by spectrally diverse communities can be explained by a positive feedback, whereby high spectral dissimilarity is both a consequence of – and results in – greater resource capture and increased investment compared to spectrally depauperate communities. Given that more spectrally dissimilar species tend to be more functionally dissimilar [13, 15], we interpret the total spectral space occupied by a plant community as a measure of its functional complexity and diversity.

### (c) Future applications of plant spectral hypervolumes

Investigating changes in the size, shape, overlap and position of spectral hypervolumes of individual plants, species and communities as they respond to a changing environment provides rich avenues for further exploration of the linkages between spectroscopy and ecological theory. Identifying the key spectral features and related anatomical, morphological and architectural characteristics which contribute most to these shifts in spectral hyperspace can indicate temporal and spatial resource partitioning [55]. Empty volumes (holes) in the spectral space occupied by plant communities might indicate colonisation or invasion potential. Alternatively, they might indicate biologically unrealised spectral types [56], because biophysical trade-offs in plant functional traits, such as between nitrogen content or photosynthetic capacity and SLA [22], limit the degree of spectral variation possible. These limitations might cause species’ spectra to diverge or converge in particular cases, and may also cause species or functional groups to follow unique trajectories in spectral space depending on environmental conditions and phenology. Further research is needed to investigate the degree to which such changes in spectral hypervolumes might be useful for detecting environmental change in its early changes and for determining the optimal timing for identifying plants spectrally. In our case, species’ hypervolume size and position in spectral and trait space changed with neighbourhood composition and community diversity (electronic supplementary material, figures S10 and S11) in parallel with species-specific trait changes with community diversity (electronic supplementary material, figure S12, tables S6 and S7). Incorporating hypervolume shifts through time in spectral plant identification models would allow plant taxa that are spectrally similar at one point in time to be differentiated with spectral time series data. Such an approach would be particularly useful in diverse environments and for large-scale studies using remote sensing data.

### (d) Conclusions

Here, we propose the measurement of spectral hypervolumes – or the N-dimensional spectral space occupied by plants – as a novel means to investigate species differentiation associated with complementarity in resource-use by individual plants and within whole communities. This method builds upon a rich history of ecological theory related to resource-use and functional diversity. We show that plants with contrasting functional attributes and evolutionary histories occupy distinct spectral spaces, providing a basis for identifying plant species, lineages and functional groups spectrally. The ecological value of spectral hypervolumes stems from the fact that plants’ reflectance spectra integrate many important dimensions of plant form and function [5, 15], including biochemical, anatomical and morphological characteristics, that are related to resource capture and investment. We found that the growth of focal plants was associated with the size of the spectral hypervolume occupied by these individuals and that ecosystem aboveground productivity was associated with the size of the spectral hypervolume occupied by a plant community. Notably, ecosystem productivity was predominantly explained by spectral complementarity—the multidimensional separation of plants in spectral space—and not by the large spectral hypervolumes occupied by productive individuals. Spectral hypervolumes unite ecological theory with the physics of light capture, demonstrating the potential to reveal plant-environment interactions and resource partitioning over large areas using rapid field optical sampling or suitable remote sensing methods. Key to implementing this approach across large spatial extents will be developing scale-appropriate sampling methods that can capture the necessary spectral information at the right temporal, spatial and spectral scales.

## Funding

This work was supported by the National Science Foundation and National Aeronautics and Space Administration through the Dimensions of Biodiversity program (DEB-1342872 grant to JCB and SEH, DEB-1342778 grant to PAT, DEB-1342827 grant to MDM., DEB-1342823 grant to JAG), by the Cedar Creek National Science Foundation Long-Term Ecological Research program (DEB-1234162), DBI 2021898 to JCB, iCORE/AITF (G224150012 and 200700172), NSERC (RGPIN-2015-05129), and CFI (26793) grants to JAG, by the University of Minnesota (Doctoral Dissertation Fellowship to SK and JJG), and by University of Minnesota’s Department of Ecology, Evolution, and Behavior (Summer Research, Crosby, Rothman, Wilkie, Anderson, and Dayton funds to JJG). AKS acknowledges the support of the Research Priority Program in Global Change and Biodiversity (URPP GCB) at the University of Zürich

## Acknowledgements

We would like to thank the following people for help in the field and lab: Brett Fredericksen for leaf-level sampling and HPLC, Ian Carriere, Erin Murdock, Ripley French and Travis Cobb for leaf-level sampling, Cathleen Lapadat for chemical assays and HPLC. Thanks to Etienne Laliberté for comments on earlier versions of the manuscript.

## Notes

### Competing Interest Statement

The authors have declared no competing interest.

### Summary of Updates

We now call the spectral space occupied by plants their "spectral hypervolume".

