## Supplementary material for "Coupling spectral and resource-use complementarity in experimental grassland and forest communities": figure S

**Figure S1.** Species clustering along linear discriminant (LD) axes maximising the differences among species based on *(a-c)* spectra and *(d-f)* foliar traits. The amount of the total variation explained by each LD axis is shown in parentheses; for species abbreviations and number of individuals per species, see figure 1.

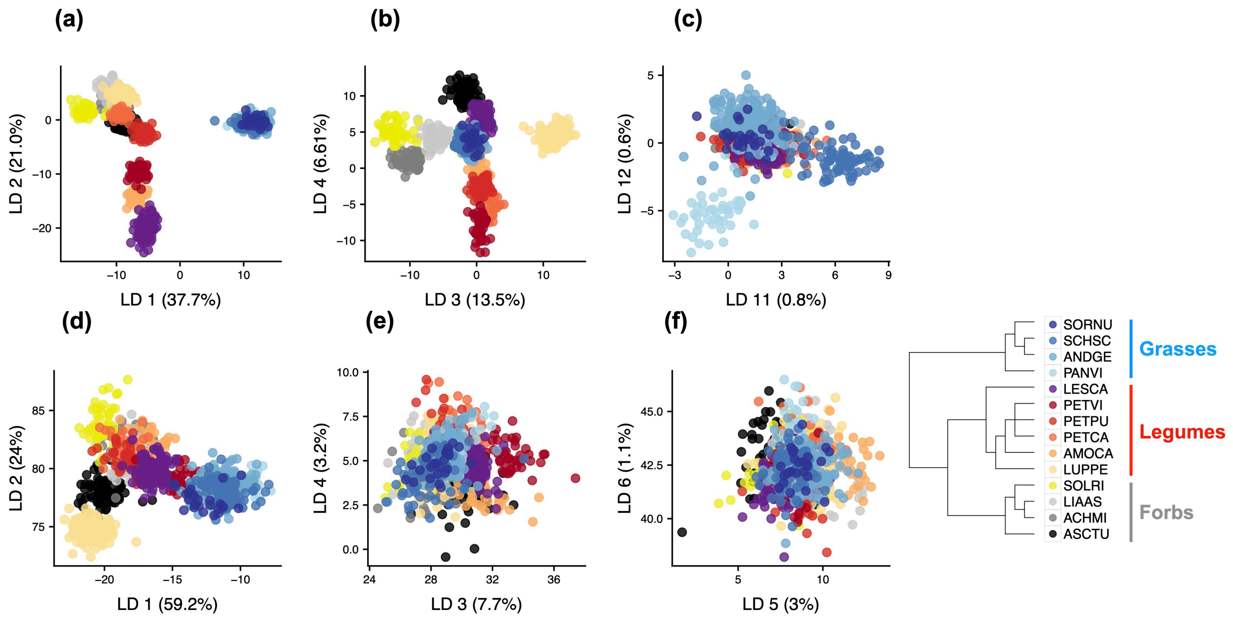

**Figure S2.** Inter- and intraspecific variation in 10 foliar traits measured in the BioDIV experiment. Measured traits are *(a)* leaf carbon, *(b)* nitrogen, *(c)* non-structural carbohydrate (NSC), *(d)* hemicellulose, *(e)* cellulose, *(f)* lignin concentration (%), *(g)* chlorophyll a+b (CHL) content (µmol $m^{-2}$), the ratios of *(h)* β-carotene (CAR) and *(i)* lutein content, and *(j)* xanthophyll pigment pool size (VAZ = violaxanthin + antheraxanthin + zeaxanthin) relative to total chlorophyll content. Dots and error bars indicate the mean +/- one standard deviation, respectively. The phylogenetic relationships among species are displayed on the left; graminoids are coded in blue, legumes in purple-red-orange and forbs in yellow-grey-black colours. For species abbreviations and number of individuals per species, see figure 1; for statistics, see electronic supplementary material, table S4.

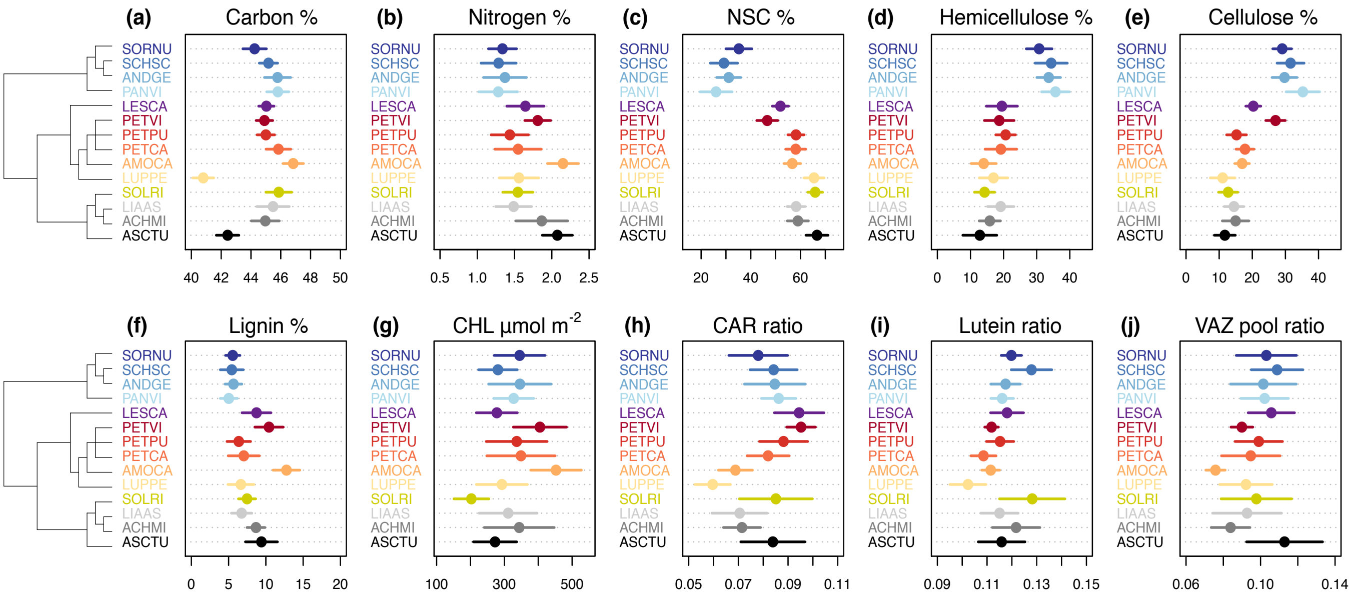

**Figure S3.** Degree of overlap among species hypervolumes in two-dimensional spectral and trait space. Illustrated are all combinations of (left) the 10 most variable spectral bands and (right) the 10 foliar traits measured in our study, respectively. Traits are centred and scaled to zero mean and a standard deviation (SD) of one (axes range from -1 to 1 SD); species means are indicated by white outlined circles; for abbreviations and number of individuals per species, see figure 1 and electronic supplementary material, figure S2.

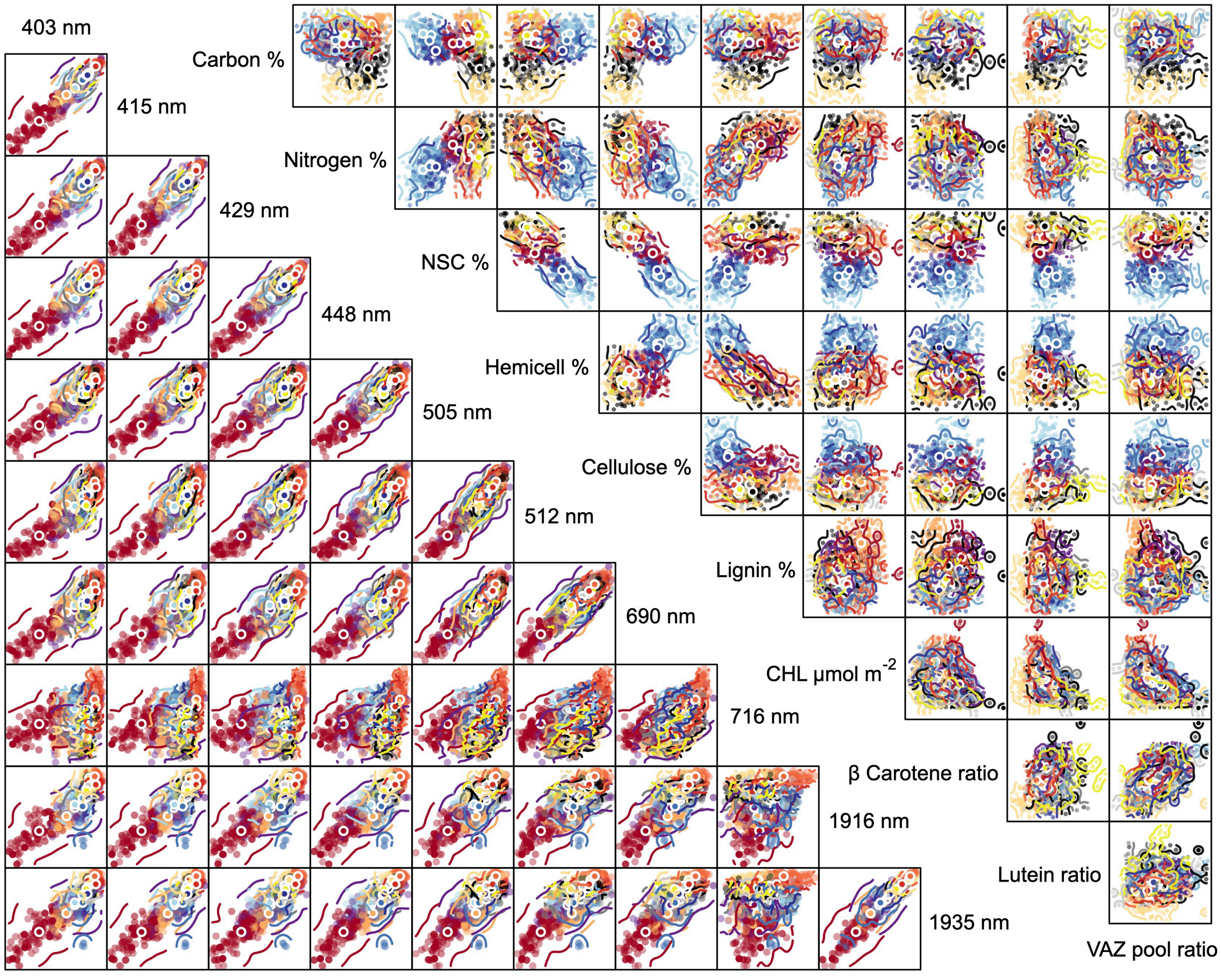

**Figure S4.** Species identification models for the BioDIV experiment. The confusion table plots for partial least squares discriminant analysis (PLSDA) models show the proportion of correctly identified (diagonal) and misidentified (off-diagonal) species based on foliar *(a)* spectra and *(b)* traits. For species abbreviations and number of individuals per species, see figure 1, for summary statistics, see electronic supplementary material, table S5.

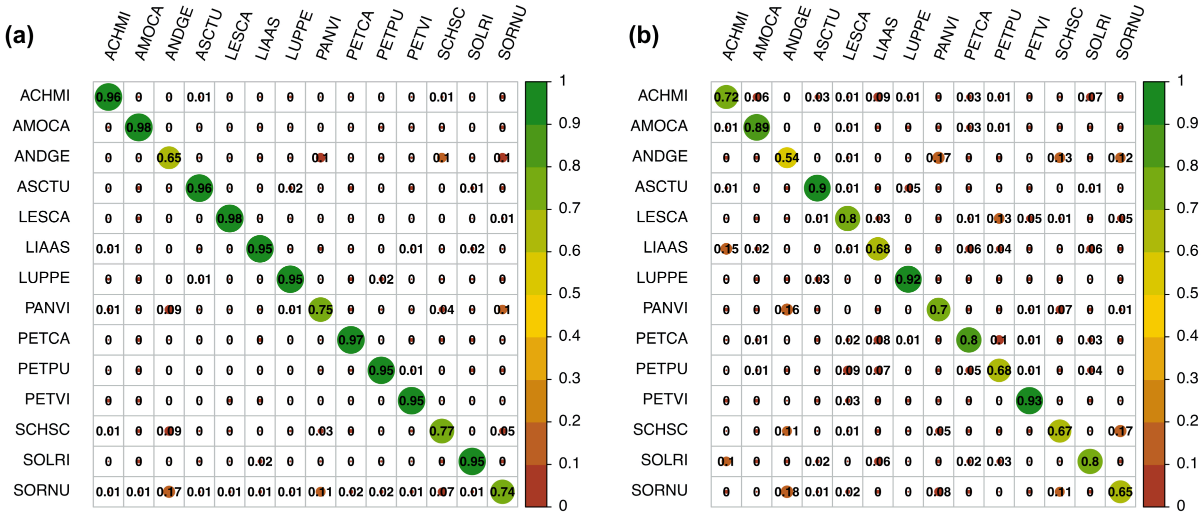

**Figure S5.** Influence of *(a)* spectral bands and *(b)* foliar traits for identifying 14 prairie-grassland perennials (n = 902) measured in the BioDIV experiment based on the absolute values of PLSDA loadings [abs(loadings), one standard deviation is indicated in grey]. For abbreviations and the number of individuals per species, see figure 1 and electronic supplementary material, figure S2.

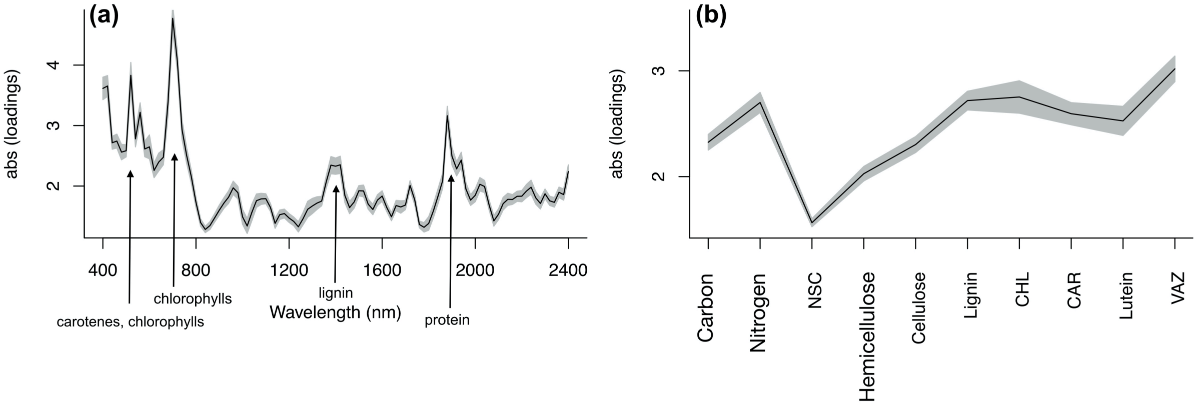

**Figure S6.** Example results for tests of phylogenetic signal (Blomberg's K). Phylogenetic signal (Blomberg's K) of the two traits showing *(a)* the strongest [hemicellulose content (%)] and *(b)* weakest [lutein ratio = lutein content (µmol $m^{-2}$) relative to total chlorophyll content (µmol $m^{-2}$)] evidence of phylogenetic signal. Significance was tested by comparing the observed K value (yellow line) to the distribution of the K statistics estimated from both a white noise (light grey bars, mean indicated by black line) and a Brownian motion null model (dark grey bars, mean indicated by red line). If observed K values are *(a)* not significantly different ($P < 0.05$) from the Brownian motion null model, they can be considered phylogenetically conserved. If observed K values are *(b)* not significantly different ($P < 0.05$) from the random expectation (white noise null model), they can be considered labile. We estimated the Brownian motion null model based on 1,000 simulations of Brownian motion evolution and the white noise model by randomly permuting traits values across the tips of the phylogeny 1,000 times (n per run = 902). For results for all other functional traits, see electronic supplementary material, table S1.

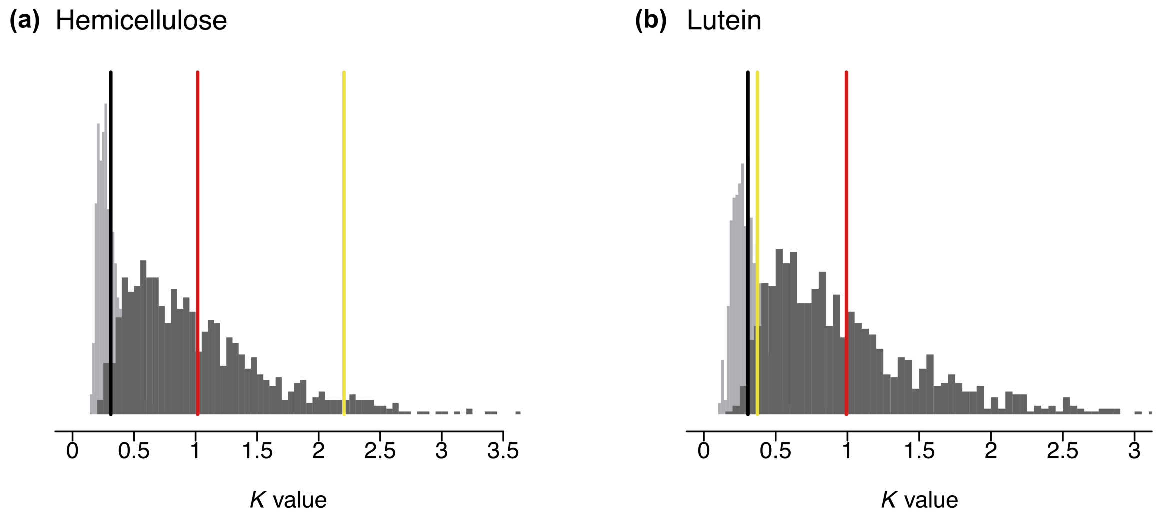

**Figure S7.** The spectral space occupied by individual trees in the FAB experiment predicting *(a)* tree height (n = 524, $r^2 = 0.14$, $F_{1,522} = 81.6$, $P < 0.001$) and *(b)* stem diameter (n = 396, $r^2 = 0.25$, $F_{1,394} = 130.2$, $P < 0.001$) in 2017. Spectral hypervolume sizes are log-transformed; the number of species per plot is indicated with different symbols.

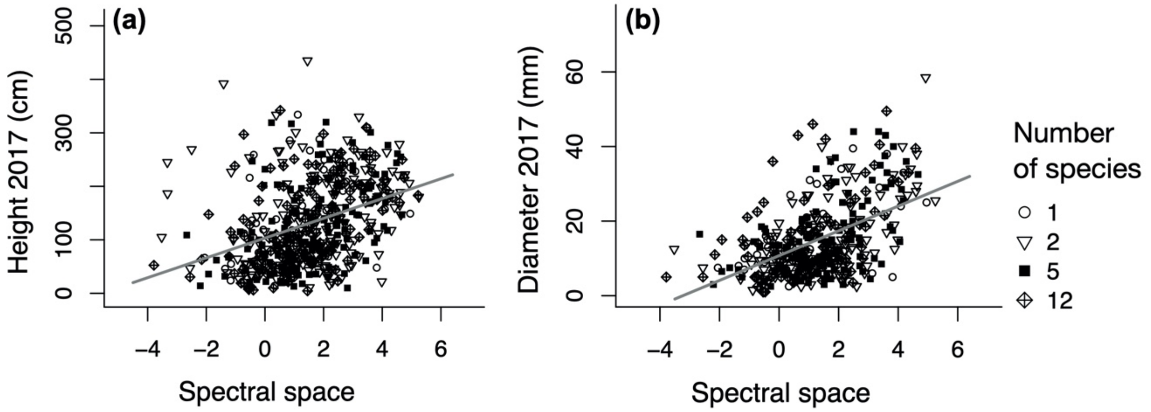


**Figure S8.** Relationship between the spectral space occupied by plant communities and species richness (the number of species per plot, indicated with different symbols) in the *(a)* BioDIV (n = 30, $r^2 = 0.25$, $F_{1,28} = 9.1$, $P < 0.001$) and *(b)* FAB experiments (n = 67, $r^2 = 0.27$, $F_{1,65} = 23.9$, $P < 0.001$); spectral hypervolume sizes are log-transformed.

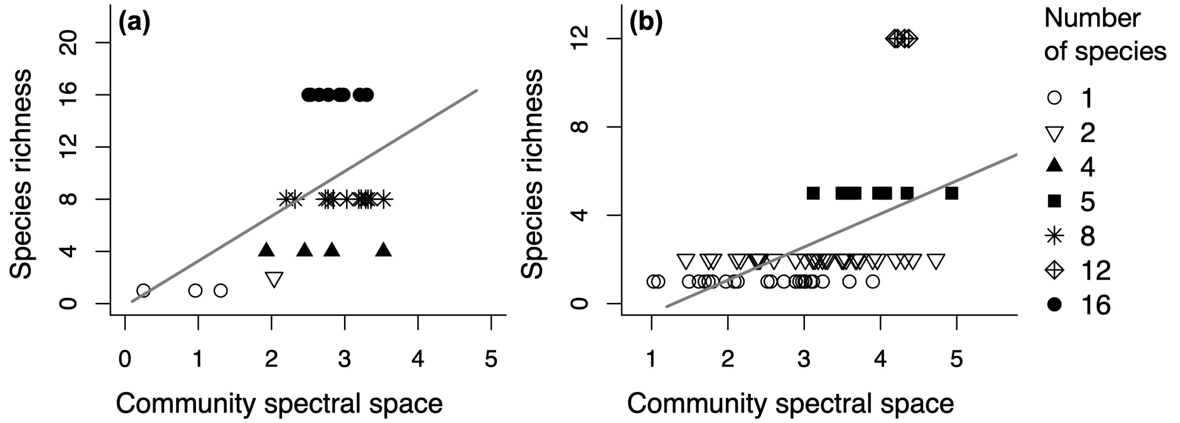

**Figure S9.** The spectral space occupied by plant communities predicting *(a)* complementarity (CE, n = 43, $r^2 = 0.36$, $F_{1,41} = 23.4$, $P < 0.001$) and *(b)* selection effect (SE, n = 43, $r^2 = 0.20$, $F_{1,41} = 10.3$, $P < 0.003$) in the FAB experiment; species richness (indicated with different symbols) is the number of species planted per plot; spectral hypervolume sizes are log-transformed.

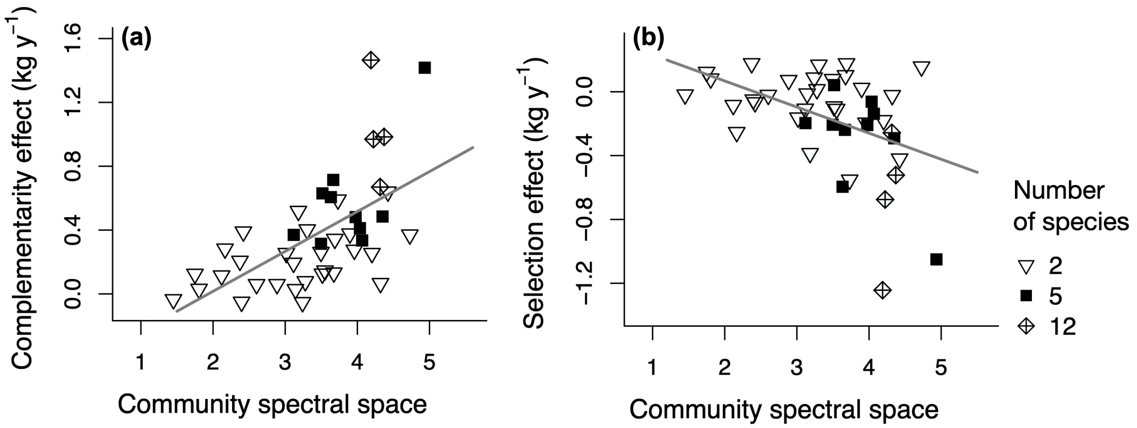


**Figure S10.** Species hypervolume shifts in trait space depending on the diversity level of the plant community. Species were measured in the BioDIV experiment; the trait space is delineated by the first three linear discriminant axes (LDs); for species abbreviations and number of individuals per species, see figure 1.

Fig S10_LDA_traits_divlevel.html

**Figure S11.** Species hypervolume shifts in spectral space depending on the diversity level of the plant community. Species were measured in the BioDIV experiment; the spectral space is delineated by the first three linear discriminant axes (LDs); for species abbreviations and number of individuals per species, see figure 1.

Fig S11_LDA_spectra_divlevel.html

**Figure S12.** Foliar trait variation depends on species identity and the diversity of the plant community. Data were measured in the BioDIV experiment; the lines represent the best linear models (linear relationships or second order polynomials) between foliar traits and species richness fitted for each species separately; when the slope of the line is nonzero, species richness influences trait expression; the 95% confidence interval is shown in grey; for details, see electronic supplementary material, appendix S1. For species abbreviations and the number of individuals per species, see figure1, for summary statistics, see electronic supplementary material, tables S4 and S7. Here, we included only species found in at least four out of the five diversity levels (1, 2, 4, 8, 16 species per plot) of the experiment.

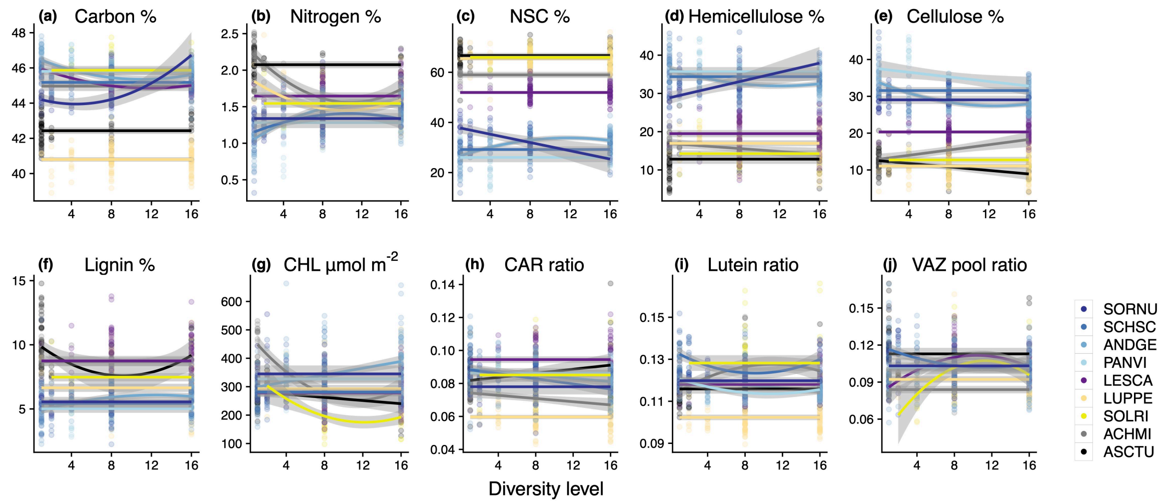
