## Supplementary material for "Coupling spectral and resource-use complementarity in experimental grassland and forest communities": table S

**Table S1.** Phylogenetic signal of 10 foliar traits measured in the BioDIV experiment (n = 902) based on Blomberg's K. Summary statistics for Blomberg's K value (K trait), the mean of the white noise model (K null mean, 1000 simulations), the mean of the Brownian motion null model (K brown mean, 1000 simulations), the number of simulations the white noise model K was greater than Blomberg's K (K null > K trait), the number of simulations the Brownian motion model K was greater than Blomberg's K (B brown > K trait), and the P values of observed vs. random variance of phylogenetic independent contrasts (PIC; P < 0.05 indicates non-random phylogenetic signal, shown in bold; see also electronic supplementary material, figure S6).

| Foliar tait | K trait | K null mean | K brown mean | K null > K trait | K brown > Ktrait | PIC P |
|---|---|---|---|---|---|---|
| Carbon % | 0.4679 | 0.3212 | 1.0088 | 129 | 864 | 0.129 |
| Nitrogen % | 0.5342 | 0.3176 | 1.0205 | 53 | 802 | **0.034** |
| Non-structural carbohydrates % | 1.6535 | 0.3030 | 1.0022 | 0 | 143 | **0.002** |
| Hemicellulose % | 2.2057 | 0.3112 | 1.0167 | 1 | 48 | **0.001** |
| Cellulose % | 1.0187 | 0.3139 | 1.0075 | 7 | 394 | **0.001** |
| Lignin % | 0.5037 | 0.3144 | 0.9822 | 73 | 819 | 0.056 |
| Chlorophyll µmol.m$^{-2}$ | 0.3885 | 0.3114 | 0.9568 | 207 | 927 | 0.235 |
| β-Carotene ratio | 0.4310 | 0.3049 | 0.9859 | 103 | 896 | 0.113 |
| Lutein ratio | 0.3724 | 0.3070 | 0.9929 | 224 | 948 | 0.228 |
| Xanthophyll pool ratio | 0.5470 | 0.3104 | 1.0005 | 38 | 777 | **0.044** |

**Table S2.** Species' unique hypervolume fractions in spectral space [mean %, (standard deviation)] with increasing number of dimension (2-21 randomly selected spectral bands); for species abbreviations and number of individuals per species see figure 1.

| | ACHMI | AMOCA | ANDGE | ASCTU | LESCA | LIAAS | LUPPE | PANVI | PETCA | PETPU | PETVI | SCHSC | SOLRI |
|---|---|---|---|---|---|---|---|---|---|---|---|---|---|
| **2 bands** | 14.1 (7.8) | 23.4 (10.9) | 6.4 (1.2) | 13.3 (6.1) | 25.3 (8.7) | 7.1 (2.2) | 33.9 (17.0) | 8.7 (3.3) | 25.1 (9.8) | 15.6 (6.2) | 54.7 (22.0) | 21.0 (10.7) | 10.9 (5.8) |
| **3 bands** | 14.4 (5.8) | 22.8 (10.6) | 6.6 (1.1) | 13.4 (5.7) | 22.4 (8.9) | 9.2 (1.4) | 33.4 (13.3) | 9.5 (1.8) | 22.0 (10.9) | 15.4 (6.3) | 60.1 (18.6) | 20.7 (9.1) | 13.7 (4.5) |
| **4 bands** | 17.8 (8.8) | 29.3 (14.1) | 6.7 (1.2) | 15.4 (6.7) | 22.0 (7.5) | 9.4 (1.5) | 34.9 (13.3) | 9.2 (1.3) | 27.6 (12.6) | 16.6 (7.8) | 64.8 (15.2) | 20.7 (8.2) | 10.2 (3.2) |
| **5 bands** | 19.0 (9.8) | 35.4 (16.1) | 5.9 (1.1) | 18.3 (8.4) | 25.2 (10.4) | 8.6 (1.7) | 41.5 (9.8) | 9.5 (2.2) | 30.8 (12.2) | 18.8 (7.4) | 66.7 (17.5) | 21.8 (7.9) | 11.0 (2.7) |
| **6 bands** | 25.8 (14.1) | 47.4 (17.3) | 8.1 (1.2) | 25.5 (12.3) | 33.8 (12.4) | 10.6 (3.6) | 50.3 (9.2) | 13.2 (2.6) | 38.0 (15.7) | 26.8 (11.2) | 78.3 (16.2) | 27.3 (8.4) | 13.6 (4.4) |
| **7 bands** | 34.9 (13.5) | 58.8 (15.2) | 8.8 (1.5) | 30.9 (11.3) | 44.4 (11.3) | 12.7 (5.1) | 58.4 (11.2) | 13.5 (6.4) | 45.1 (16.0) | 30.9 (10.8) | 89.4 (7.0) | 33.3 (9.5) | 15.5 (7.0) |
| **8 bands** | 48.0 (18.6) | 70.5 (14.8) | 11.2 (3.9) | 41.9 (16.5) | 57.1 (11.9) | 19.3 (9.3) | 67.5 (9.6) | 18.1 (9.0) | 57.0 (18.3) | 42.8 (15.7) | 94.2 (4.9) | 43.6 (10.2) | 28.0 (12.9) |
| **9 bands** | 58.3 (16.7) | 81.2 (10.5) | 15.6 (8.2) | 54.5 (13.6) | 67.0 (11.1) | 27.8 (12.0) | 76.7 (8.3) | 25.5 (12.2) | 68.9 (12.7) | 55.7 (20.2) | 97.1 (2.9) | 54.3 (9.2) | 33.4 (14.6) |
| **10 bands** | 75.9 (9.9) | 90.5 (5.0) | 25.2 (12.4) | 70.0 (9.0) | 80.2 (6.9) | 45.2 (15.6) | 87.2 (5.4) | 39.9 (11.0) | 80.4 (12.4) | 68.9 (16.4) | 98.7 (1.6) | 67.0 (6.6) | 55.7 (19.4) |
| **11 bands** | 84.0 (8.4) | 95.6 (3.0) | 44.1 (17.5) | 80.8 (9.7) | 88.3 (5.8) | 62.9 (17.2) | 93.3 (3.8) | 53.7 (11.9) | 88.6 (9.2) | 85.1 (12.9) | 99.6 (0.7) | 77.2 (5.4) | 72.8 (17.1) |
| **12 bands** | 91.6 (6.0) | 97.3 (2.4) | 59.9 (22.5) | 90.1 (6.4) | 92.9 (3.8) | 80.9 (17.1) | 96.7 (3.3) | 68.2 (13.3) | 94.6 (6.8) | 93.6 (6.3) | 99.8 (0.4) | 85.4 (5.2) | 83.8 (16.9) |
| **13 bands** | 95.3 (4.8) | 98.8 (1.0) | 70.6 (21.0) | 94.0 (5.3) | 95.4 (2.9) | 85.8 (20.7) | 97.9 (3.4) | 79.9 (10.9) | 97.2 (3.4) | 95.2 (10.4) | 99.9 (0.2) | 90.9 (3.5) | 89.3 (13.2) |
| **14 bands** | 97.7 (3.4) | 99.4 (0.9) | 87.7 (13.7) | 96.7 (4.2) | 98.0 (1.7) | 92.2 (14.6) | 98.8 (3.2) | 84.8 (13.5) | 97.9 (3.4) | 99.2 (1.7) | 100.0 (0.1) | 95.1 (2.8) | 91.6 (16.2) |
| **15 bands** | 99.3 (1.6) | 99.9 (0.2) | 89.6 (14.0) | 98.6 (2.5) | 99.2 (0.8) | 98.4 (8.6) | 99.8 (0.8) | 95.2 (4.8) | 99.8 (0.9) | 99.4 (1.2) | 100.0 (0.0) | 98.0 (0.9) | 96.7 (12.8) |
| **16 bands** | 99.5 (1.4) | 99.9 (0.3) | 100.0 (1.0) | 99.6 (1.3) | 99.6 (0.4) | 99.3 (2.6) | 99.8 (0.8) | 97.3 (5.6) | 98.9 (5.5) | 98.1 (6.7) | 100.0 (0.0) | 98.7 (1.1) | 99.1 (2.5) |
| **17 bands** | 100.0 (0.2) | 100.0 (0.1) | 100.0 (1.3) | 99.8 (0.6) | 99.9 (0.2) | 100.1 (1.0) | 99.7 (2.0) | 98.4 (4.6) | 100.0 (0.3) | 100.1 (0.8) | 100.0 (0.0) | 99.3 (0.6) | 100.0 (1.0) |
| **18 bands** | 100.0 (0.1) | 100.0 (0.1) | 100.0 (1.0) | 100.0 (0.3) | 100.0 (0.1) | 99.0 (2.4) | 100.0 (0.2) | 99.1 (3.8) | 100.0 (0.3) | 100.1 (0.8) | 100.0 (0.0) | 99.8 (0.3) | 100.1 (1.1) |
| **19 bands** | 100.0 (0.0) | 100.0 (0.0) | 100.0 (1.2) | 99.8 (1.0) | 100.0 (0.1) | 100.1 (1.0) | 100.0 (0.2) | 99.5 (2.4) | 100.0 (0.1) | 100.1 (0.9) | 100.0 (0.0) | 99.8 (0.9) | 99.3 (1.3) |
| **20 bands** | 99.9 (0.3) | 100.0 (0.0) | 100.1 (1.2) | 99.9 (0.3) | 100.0 (0.1) | 99.6 (1.1) | 100.0 (0.0) | 99.9 (0.6) | 99.9 (0.3) | 99.7 (0.8) | 100.0 (0.0) | 99.9 (0.3) | 100.1 (1.1) |
| **21 bands** | 100.0 (0.1) | 100.0 (0.0) | 100.0 (1.2) | 100.0 (0.2) | 100.0 (0.0) | 100.2 (1.1) | 100.0 (0.3) | 99.9 (0.4) | 100.0 (0.3) | 100.0 (0.9) | 100.0 (0.0) | 100.0 (0.0) | 99.6 (1.0) |

**Table S3.** Species 'unique hypervolume fractions in trait space [mean %, (standard deviation)] with increasing number of dimension (2-10 randomly selected foliar traits); for species abbreviations and number of individuals per species see figure 1.

| | ACHMI | AMOCA | ANDGE | ASCTU | LESCA | LIAAS | LUPPE | PANVI | PETCA | PETPU | PETVI | SCHSC | SOLRI | SORNU |
|---|---|---|---|---|---|---|---|---|---|---|---|---|---|---|
| **2 traits** | 14.0 (8.8) | 29.3 (20.6) | 18.1 (9.5) | 36.3 (15.5) | 8.5 (4.6) | 12.7 (7.5) | 35.2 (28.8) | 19.5 (12.4) | 11.4 (6.5) | 7.5 (4.1) | 14.4 (12.3) | 21.2 (11.1) | 17.8 (13.4) | 10.1 (5.4) |
| **3 traits** | 11.2 (7.6) | 31.9 (22.0) | 16.7 (8.5) | 41.7 (18.6) | 8.4 (5.8) | 11.1 (5.5) | 52.9 (30.6) | 11.6 (8.4) | 10.2 (5.5) | 6.4 (4.3) | 12.6 (10.7) | 17.0 (10.4) | 21.4 (13.5) | 9.1 (5.8) |
| **4 traits** | 14.8 (9.2) | 38.1 (21.9) | 17.7 (6.9) | 45.6 (17.8) | 12.4 (6.3) | 11.7 (6.1) | 54.6 (25.8) | 14.0 (7.2) | 12.9 (5.7) | 8.0 (5.1) | 26.8 (15.0) | 20.6 (11.4) | 22.5 (11.8) | 10.5 (5.5) |
| **5 traits** | 21.4 (8.7) | 50.5 (23.9) | 21.0 (6.5) | 63.4 (16.4) | 13.9 (7.2) | 19.1 (9.2) | 70.2 (25.4) | 16.9 (8.9) | 19.4 (6.8) | 11.1 (5.4) | 28.4 (13.2) | 20.6 (10.7) | 31.7 (14.8) | 14.1 (7.4) |
| **6 traits** | 40.4 (11.9) | 63.9 (20.7) | 31.0 (7.7) | 75.7 (14.3) | 26.2 (7.7) | 32.3 (9.9) | 82.4 (18.4) | 24.8 (9.5) | 36.9 (9.6) | 22.2 (9.5) | 46.4 (16.6) | 32.5 (11.8) | 49.3 (15.0) | 24.2 (8.4) |
| **7 traits** | 58.2 (12.2) | 77.9 (18.7) | 44.0 (6.9) | 85.6 (12.3) | 40.6 (9.8) | 50.2 (11.9) | 91.3 (11.1) | 36.6 (7.1) | 55.1 (9.7) | 37.4 (8.7) | 69.1 (14.2) | 43.5 (10.5) | 68.6 (11.2) | 40.3 (9.2) |
| **8 traits** | 77.3 (7.8) | 91.4 (6.8) | 58.5 (5.9) | 97.3 (3.0) | 60.7 (7.1) | 70.9 (8.0) | 98.6 (1.7) | 50.0 (6.6) | 73.9 (7.6) | 57.9 (9.8) | 82.9 (7.9) | 58.5 (8.9) | 85.2 (6.8) | 59.8 (8.4) |
| **9 traits** | 90.3 (3.2) | 96.9 (3.7) | 74.7 (4.8) | 99.3 (0.7) | 80.7 (5.9) | 84.5 (5.2) | 99.7 (0.5) | 68.4 (4.6) | 88.0 (3.6) | 75.7 (6.7) | 94.0 (6.0) | 74.6 (4.9) | 94.2 (2.7) | 76.7 (5.0) |
| **10 traits** | 96.8 (1.0) | 99.3 (1.3) | 86.7 (3.5) | 99.9 (0.1) | 92.4 (4.3) | 94.4 (1.4) | 100.0 (0.1) | 83.2 (2.6) | 95.2 (1.6) | 89.8 (3.6) | 98.0 (2.8) | 86.1 (2.3) | 98.2 (1.0) | 89.2 (2.2) |

**Table S4.** Differences in foliar traits among species. Data were collected in the BioDIV experiment; n is the number of species; significant differences ($P < 0.05$) based on Tukey's HSD test among species are indicated with letters; for species abbreviations see figure 1.

| Foliar trait | Species | Mean (SD) | Range | Tukey |
|---|---|---|---|---|
| **Carbon (%)** | SORNU | 44.24 (0.15) | 43.22, 46.74 | e |
| | SCHSC | 45.18 (0.07) | 43.46, 46.44 | cd |
| | ANDGE | 45.78 (0.07) | 43.14, 47.81 | b |
| | PANVI | 45.80 (0.11) | 43.78, 47.33 | b |
| | LESCA | 45.03 (0.05) | 43.89, 46.78 | cd |
| | PETVI | 44.91 (0.09) | 43.80, 46.34 | d |
| | PETPU | 45.00 (0.08) | 43.81, 46.29 | cd |
| | PETCA | 45.84 (0.16) | 44.31, 47.67 | b |
| | AMOCA | 46.84 (0.13) | 45.38, 47.88 | a |
| | LUPPE | 40.80 (0.06) | 38.90, 42.32 | g |
| | SOLRI | 45.87 (0.12) | 43.18, 47.74 | b |
| | LIAAS | 45.48 (0.16) | 43.15, 48.18 | bc |
| | ACHMI | 44.96 (0.13) | 41.24, 46.51 | cd |
| | ASCTU | 42.43 (0.09) | 40.79, 44.25 | f |
| **Nitrogen (%)** | SORNU | 1.34 (0.04) | 0.97, 1.87 | ef |
| | SCHSC | 1.29 (0.03) | 0.32, 1.80 | f |
| | ANDGE | 1.37 (0.02) | 0.60, 2.12 | ef |
| | PANVI | 1.28 (0.04) | 0.54, 1.77 | f |
| | LESCA | 1.65 (0.03) | 1.09, 2.28 | c |
| | PETVI | 1.81 (0.03) | 1.49, 2.23 | b |
| | PETPU | 1.44 (0.03) | 0.66, 1.96 | def |
| | PETCA | 1.55 (0.06) | 0.72, 2.22 | cd |
| | AMOCA | 2.15 (0.04) | 1.67, 2.49 | a |
| | LUPPE | 1.56 (0.02) | 0.94, 2.48 | cd |
| | SOLRI | 1.54 (0.03) | 1.21, 1.98 | cd |
| | LIAAS | 1.49 (0.03) | 0.86, 1.99 | de |
| | ACHMI | 1.86 (0.05) | 1.13, 2.52 | b |
| | ASCTU | 2.07 (0.02) | 1.61, 2.50 | a |
| **Nonstructural carbohydrates (%)** | SORNU | 35.21 (1.01) | 25.85, 44.70 | e |
| | SCHSC | 29.19 (0.63) | 11.90, 40.78 | f |
| | ANDGE | 31.13 (0.38) | 16.81, 42.70 | f |
| | PANVI | 26.02 (0.92) | 13.06, 40.37 | g |
| | LESCA | 51.94 (0.33) | 45.13, 60.38 | c |
| | PETVI | 46.63 (0.65) | 34.36, 58.82 | d |
| | PETPU | 58.21 (0.45) | 50.26, 65.03 | b |
| | PETCA | 58.09 (0.76) | 50.85, 65.29 | b |
| | AMOCA | 56.72 (0.64) | 51.14, 63.65 | b |
| | LUPPE | 65.46 (0.37) | 49.06, 76.02 | a |
| | SOLRI | 65.93 (0.43) | 59.77, 72.55 | a |
| | LIAAS | 58.26 (0.52) | 50.02, 67.95 | b |
| | ACHMI | 58.96 (0.61) | 49.33, 68.81 | b |
| | ASCTU | 66.66 (0.54) | 48.73, 74.95 | a |

| | | | | |
|---|---|---|---|---|
| **Hemicellulose (%)** | SORNU | 30.74 (0.75) | 21.48, 38.59 | b |
| | SCHSC | 34.38 (0.56) | 15.42, 46.03 | a |
| | ANDGE | 33.62 (0.28) | 25.00, 44.65 | ab |
| | PANVI | 35.67 (0.60) | 27.48, 44.23 | a |
| | LESCA | 19.48 (0.47) | 7.33, 29.30 | c |
| | PETCA | 18.69 (0.69) | 9.71, 29.62 | cde |
| | AMOCA | 20.61 (0.42) | 13.43, 26.45 | c |
| | LUPPE | 19.19 (0.91) | 12.36, 30.07 | cd |
| | SOLRI | 14.00 (0.72) | 3.97, 20.30 | fg |
| | LIAAS | 16.96 (0.39) | 5.91, 30.24 | de |
| | ACHMI | 14.26 (0.44) | 8.73, 22.83 | fg |
| | ASCTU | 19.13 (0.57) | 11.16, 28.38 | cd |
| | BERIN | 15.83 (0.46) | 8.90, 22.51 | ef |
| | RUMAC | 12.82 (0.61) | 4.00, 28.06 | g |
| **Cellulose (%)** | SORNU | 29.04 (0.55) | 24.02, 34.43 | cd |
| | SCHSC | 31.56 (0.47) | 22.07, 47.44 | b |
| | ANDGE | 29.74 (0.30) | 23.68, 43.70 | c |
| | PANVI | 35.27 (0.72) | 25.75, 46.14 | a |
| | LESCA | 20.32 (0.23) | 14.39, 27.69 | e |
| | PETVI | 27.02 (0.46) | 21.71, 33.40 | d |
| | PETPU | 15.25 (0.42) | 7.83, 22.58 | fg |
| | PETCA | 17.80 (0.51) | 12.39, 23.24 | f |
| | AMOCA | 16.95 (0.42) | 13.18, 21.86 | fg |
| | LUPPE | 11.06 (0.34) | 3.77, 22.20 | i |
| | SOLRI | 12.73 (0.41) | 5.67, 18.92 | hi |
| | LIAAS | 14.48 (0.43) | 5.00, 21.04 | gh |
| | ACHMI | 14.96 (0.57) | 7.46, 22.49 | gh |
| | ASCTU | 11.72 (0.38) | 4.04, 20.69 | i |
| **Lignin (%)** | SORNU | 5.55 (0.19) | 3.45, 7.66 | fg |
| | SCHSC | 5.43 (0.18) | 2.25, 11.98 | fg |
| | ANDGE | 5.65 (0.09) | 2.27, 8.79 | fg |
| | PANVI | 5.03 (0.16) | 2.83, 7.14 | g |
| | LESCA | 8.75 (0.20) | 4.70, 13.78 | c |
| | PETVI | 10.42 (0.30) | 6.81, 15.05 | b |
| | PETPU | 6.37 (0.22) | 2.39, 9.92 | ef |
| | PETCA | 7.04 (0.40) | 2.86, 10.02 | de |
| | AMOCA | 12.77 (0.34) | 9.33, 16.74 | a |
| | LUPPE | 6.65 (0.16) | 2.89, 11.76 | de |
| | SOLRI | 7.48 (0.16) | 5.49, 10.22 | d |
| | LIAAS | 6.75 (0.20) | 3.80, 9.71 | de |
| | ACHMI | 8.68 (0.17) | 5.73, 10.66 | c |
| | ASCTU | 9.41 (0.26) | 4.66, 14.78 | bc |
| **Chlorophyll a+b (µmol $m^{-2}$)** | SORNU | 345.11 (14.49) | 250.82, 555.24 | cd |
| | SCHSC | 280.93 (6.58) | 183.78, 495.04 | ef |
| | ANDGE | 345.96 (7.27) | 158.37, 663.35 | c |
| | PANVI | 327.49 (8.46) | 216.02, 508.41 | cde |
| | LESCA | 277.76 (6.10) | 150.50, 437.34 | f |
| | PETVI | 404.99 (12.12) | 281.97, 682.72 | ab |
| | PETPU | 336.42 (12.49) | 198.82, 630.79 | cd |

| | | | | |
|---|---|---|---|---|
| | PETCA | 349.04 (19.11) | 126.10, 556.47 | bc |
| | AMOCA | 452.52 (14.17) | 304.00, 598.69 | a |
| | LUPPE | 292.70 (6.87) | 124.72, 503.96 | def |
| | SOLRI | 202.62 (7.31) | 97.41, 329.47 | g |
| | LIAAS | 311.14 (12.22) | 144.52, 515.50 | cdef |
| | ACHMI | 343.87 (14.76) | 177.83, 531.39 | cd |
| | ASCTU | 272.49 (7.64) | 126.94, 414.50 | f |
| **β-Carotene ratio (μmol m$^{-2}$)** | SORNU | 0.08 (0.00) | 0.05, 0.10 | cd |
| | SCHSC | 0.08 (0.00) | 0.06, 0.10 | bc |
| | ANDGE | 0.08 (0.00) | 0.05, 0.12 | bc |
| | PANVI | 0.09 (0.00) | 0.08, 0.10 | bc |
| | LESCA | 0.09 (0.00) | 0.07, 0.12 | a |
| | PETVI | 0.10 (0.00) | 0.09, 0.11 | a |
| | PETPU | 0.09 (0.00) | 0.07, 0.12 | ab |
| | PETCA | 0.08 (0.00) | 0.07, 0.10 | bc |
| | AMOCA | 0.07 (0.00) | 0.05, 0.08 | d |
| | LUPPE | 0.06 (0.00) | 0.04, 0.08 | e |
| | SOLRI | 0.09 (0.00) | 0.06, 0.12 | bc |
| | LIAAS | 0.07 (0.00) | 0.05, 0.11 | d |
| | ACHMI | 0.07 (0.00) | 0.05, 0.09 | d |
| | ASCTU | 0.08 (0.00) | 0.05, 0.14 | bc |
| **Lutein ratio (μmol m$^{-2}$)** | SORNU | 0.12 (0.00) | 0.11, 0.13 | bc |
| | SCHSC | 0.13 (0.00) | 0.11, 0.15 | a |
| | ANDGE | 0.12 (0.00) | 0.11, 0.14 | c |
| | PANVI | 0.12 (0.00) | 0.11, 0.13 | cd |
| | LESCA | 0.12 (0.00) | 0.11, 0.14 | bc |
| | PETVI | 0.11 (0.00) | 0.11, 0.12 | de |
| | PETPU | 0.12 (0.00) | 0.11, 0.13 | cd |
| | PETCA | 0.11 (0.00) | 0.10, 0.12 | e |
| | AMOCA | 0.11 (0.00) | 0.11, 0.12 | de |
| | LUPPE | 0.10 (0.00) | 0.09, 0.13 | f |
| | SOLRI | 0.13 (0.00) | 0.11, 0.17 | a |
| | LIAAS | 0.12 (0.00) | 0.10, 0.14 | cd |
| | ACHMI | 0.12 (0.00) | 0.11, 0.15 | b |
| | ASCTU | 0.12 (0.00) | 0.10, 0.15 | cd |
| **VAZ pool ratio (μmol m$^{-2}$)** | SORNU | 0.10 (0.00) | 0.07, 0.13 | abcd |
| | SCHSC | 0.11 (0.00) | 0.06, 0.15 | ab |
| | ANDGE | 0.10 (0.00) | 0.06, 0.17 | cd |
| | PANVI | 0.10 (0.00) | 0.08, 0.12 | bcd |
| | LESCA | 0.11 (0.00) | 0.08, 0.14 | abc |
| | PETVI | 0.09 (0.00) | 0.08, 0.11 | ef |
| | PETPU | 0.10 (0.00) | 0.08, 0.14 | cde |
| | PETCA | 0.09 (0.00) | 0.07, 0.14 | def |
| | AMOCA | 0.08 (0.00) | 0.06, 0.09 | g |
| | LUPPE | 0.09 (0.00) | 0.06, 0.16 | ef |
| | SOLRI | 0.10 (0.00) | 0.06, 0.15 | cde |
| | LIAAS | 0.09 (0.00) | 0.06, 0.16 | def |
| | ACHMI | 0.08 (0.00) | 0.07, 0.12 | fg |
| | ASCTU | 0.11 (0.00) | 0.08, 0.17 | a |

**Table S5.** Partial least squares discriminate analysis (PLSDA) model identifying plant species in the BioDIV experiment based on spectra and foliar traits (n=902), respectively; summary statistics are the number of components in PLSDA, the mean overall classification accuracy, and the mean Kappa statistic; SD indicates the standard deviation after 100 model iterations.

| PLSDA BioDIV | summary statistics | spectra | traits | training samples |
|---|---|---|---|---|
| | components | 25 | 9 | 20 |
| | accuracy (SD) | 0.93 (0.01) | 0.66 (0.02) | |
| | Kappa (SD) | 0.81 (0.02) | 0.63 (0.02) | |

**Table S6.** Linear models testing the effect of species richness on foliar trait variation. Using data collected in the BioDIV experiment we tested six alternative hypothesis: (1) traits do not differ among species and do not change with increasing species richness (formula: trait ~ 1), (2) traits change with increasing species richness but do not differ among species (formula: trait ~ species richness), (3) traits differ among species but do not change with increasing species richness (formula: trait ~ species), (4) traits differ among species but changes with species richness are consistent (formula: trait ~ species + species richness), (5) traits do not differ among species in monoculture but vary among species with increasing species richness (formula: trait ~ species richness/species), (6) traits differ among species and change for each species differently with species richness (formula: trait ~ species richness*species). The best models for each foliar trait based on Akaike's information criterion (AIC) are indicated in bold. Here we included only species found in at least four out of the five diversity levels of the experiment (n = 703).

| Functional trait | Model | Formula | AIC | $r^2$ | F statistic | df | P value |
|---|---|---|---|---|---|---|---|
| Carbon % | 1 | C~1 | 2983.64 | 0 | 44.31 | 1 | <0.001 |
| Carbon % | 2 | C~species | 1624.16 | 0.86 | 527.02 | 9 | <0.001 |
| Carbon % | 3 | C~species richness | 2983.67 | 0 | 1.97 | 2 | 0.1614 |
| Carbon % | 4 | C~species richness+species | 1615.09 | 0.86 | 476.43 | 10 | <0.001 |
| Carbon % | 5 | C~species richness/species | 2334.24 | 0.61 | 121.97 | 10 | <0.001 |
| Carbon % | 6 | C~species richness*species | **1609.54** | 0.87 | 258.33 | 18 | <0.001 |
| Nitrogen % | 1 | N~1 | 533.16 | 0 | 1.54 | 1 | <0.001 |
| Nitrogen % | 2 | N~species | 116.23 | 0.46 | 73.84 | 9 | <0.001 |
| Nitrogen % | 3 | N~species richness | 533.93 | 0 | 1.23 | 2 | 0.2686 |
| Nitrogen % | 4 | N~species richness+species | 101.54 | 0.47 | 68.97 | 10 | <0.001 |
| Nitrogen % | 5 | N~species richness/species | 438.13 | 0.15 | 13.43 | 10 | <0.001 |
| Nitrogen % | 6 | N~species richness*species | **25.11** | 0.54 | 46.82 | 18 | <0.001 |
| Non-structural carbohydrates % | 1 | NSC~1 | 5972.5 | 0 | 47.51 | 1 | <0.001 |
| Non-structural carbohydrates % | 2 | NSC~species | 4148.28 | 0.93 | 1102.04 | 9 | <0.001 |
| Non-structural carbohydrates % | 3 | NSC~species richness | 5969.77 | 0.01 | 4.73 | 2 | 0.03 |
| Non-structural carbohydrates % | 4 | NSC~species richness+species | 4143.41 | 0.93 | 988.55 | 10 | <0.001 |
| Non-structural carbohydrates % | 5 | NSC~species richness/species | 5328.07 | 0.61 | 120.57 | 10 | <0.001 |
| Non-structural carbohydrates % | 6 | NSC~species richness*species | **4112.59** | 0.93 | 555.71 | 18 | <0.001 |
| Hemicellulose % | 1 | Hemicell~1 | 5214.67 | 0 | 24.19 | 1 | <0.001 |
| Hemicellulose % | 2 | Hemicell~species | 4027.92 | 0.82 | 393.31 | 9 | <0.001 |
| Hemicellulose % | 3 | Hemicell~species richness | 5212.07 | 0.01 | 4.6 | 2 | 0.0323 |
| Hemicellulose % | 4 | Hemicell~species richness+species | 4023.37 | 0.82 | 353.09 | 10 | <0.001 |
| Hemicellulose % | 5 | Hemicell~species richness/species | 4700.33 | 0.53 | 87.19 | 10 | <0.001 |

| | | | | | | | |
|---|---|---|---|---|---|---|---|
| Hemicellulose % | 6 | Hemicell~species richness*species | **4016.06** | 0.83 | 192.36 | 18 | <0.001 |
| Cellulose % | 1 | Cellulose~1 | 5176.4 | 0 | 21.72 | 1 | <0.001 |
| Cellulose % | 2 | Cellulose~species | 3814.04 | 0.86 | 529.54 | 9 | <0.001 |
| Cellulose % | 3 | Cellulose~species richness | 5169.14 | 0.01 | 9.29 | 2 | 0.0024 |
| Cellulose % | 4 | Cellulose~species richness+species | 3796.26 | 0.86 | 485.63 | 10 | <0.001 |
| Cellulose % | 5 | Cellulose~species richness/species | 4649.62 | 0.54 | 90.12 | 10 | <0.001 |
| Cellulose % | 6 | Cellulose~species richness*species | **3736.06** | 0.88 | 287.84 | 18 | <0.001 |
| Lignin % | 1 | Lignin~1 | 3078.31 | 0 | 6.9 | 1 | <0.001 |
| Lignin % | 2 | Lignin~species | 2623.7 | 0.49 | 82.68 | 9 | <0.001 |
| Lignin % | 3 | Lignin~species richness | **3079.92** | 0 | 0.39 | 2 | 0.5341 |
| Lignin % | 4 | Lignin~species richness+species | **2624.65** | 0.49 | 73.62 | 10 | <0.001 |
| Lignin % | 5 | Lignin~species richness/species | 2875.05 | 0.27 | 28.48 | 10 | <0.001 |
| Lignin % | 6 | Lignin~species richness*species | **2626.8** | 0.5 | 40.09 | 18 | <0.001 |
| Chlorophyll μmol.m$^{-2}$ | 1 | Chl~1 | 8241.33 | 0 | 301.18 | 1 | <0.001 |
| Chlorophyll μmol.m$^{-2}$ | 2 | Chl~species | 8077.88 | 0.23 | 25.23 | 9 | <0.001 |
| Chlorophyll μmol.m$^{-2}$ | 3 | Chl~species richness | 8243.08 | 0 | 0.26 | 2 | 0.6112 |
| Chlorophyll μmol.m$^{-2}$ | 4 | Chl~species richness+species | 8077.52 | 0.23 | 22.73 | 10 | <0.001 |
| Chlorophyll μmol.m$^{-2}$ | 5 | Chl~species richness/species | 8075.99 | 0.23 | 22.94 | 10 | <0.001 |
| Chlorophyll μmol.m$^{-2}$ | 6 | Chl~species richness*species | **8011.99** | 0.31 | 18.31 | 18 | <0.001 |
| β-Carotene ratio | 1 | bCar~1 | -3892.94 | 0 | 0.08 | 1 | <0.001 |
| β-Carotene ratio | 2 | bCar~species | -4388.57 | 0.52 | 92.86 | 9 | <0.001 |
| β-Carotene ratio | 3 | bCar~species richness | -3897.68 | 0.01 | 6.76 | 2 | 0.0095 |
| β-Carotene ratio | 4 | bCar~species richness+species | -4398.38 | 0.53 | 85.13 | 10 | <0.001 |
| β-Carotene ratio | 5 | bCar~species richness/species | -4271.23 | 0.43 | 58.3 | 10 | <0.001 |
| β-Carotene ratio | 6 | bCar~species richness*species | **-4407.34** | 0.54 | 47.61 | 18 | <0.001 |
| Lutein ratio | 1 | Lutein~1 | -4343.29 | 0 | 0.12 | 1 | <0.001 |
| Lutein ratio | 2 | Lutein~species | -4822.23 | 0.51 | 88.65 | 9 | <0.001 |
| Lutein ratio | 3 | Lutein~species richness | -4352.25 | 0.02 | 11.01 | 2 | <0.001 |
| Lutein ratio | 4 | Lutein~species richness+species | -4821 | 0.51 | 78.86 | 10 | <0.001 |
| Lutein ratio | 5 | Lutein~species richness/species | -4705.28 | 0.42 | 55.2 | 10 | <0.001 |
| Lutein ratio | 6 | Lutein~species richness*species | **-4842.24** | 0.53 | 45.7 | 18 | <0.001 |
| Xanthophyll pool ratio | 1 | VAZ~1 | -3701.48 | 0 | 0.1 | 1 | <0.001 |
| Xanthophyll pool ratio | 2 | VAZ~species | -3838.17 | 0.2 | 21.04 | 9 | <0.001 |
| Xanthophyll pool ratio | 3 | VAZ~species richness | -3712.5 | 0.02 | 13.11 | 2 | 0.0003 |
| Xanthophyll pool ratio | 4 | VAZ~species richness+species | -3841.42 | 0.2 | 19.4 | 10 | <0.001 |
| Xanthophyll pool ratio | 5 | VAZ~species richness/species | -3772.74 | 0.12 | 10.42 | 10 | <0.001 |
| Xanthophyll pool ratio | 6 | VAZ~species richness*species | **-3836.54** | 0.21 | 10.95 | 18 | <0.001 |

**Table S7.** Best linear models predicting the effect of species richness on foliar trait variation in the BioDIV experiment. Best models were selected based on Akaike's information criterion (AIC); models are fit for each species separately; for species abbreviations and number of individuals per species see figure 1. Here we included only species found in at least four out of the five diversity levels of the experiment, n is the number of individuals per species.

| Species | n | Foliar trait | Formula | $r^2$ | F stat. | df | P value | AIC |
|---|---|---|---|---|---|---|---|---|
| ACHMI | 49 | Carbon % | C~1 | NA | NA | 1 | NA | 136.09 |
| ANDGE | 162 | Carbon % | C~poly(species richness, 2) | 0.19 | 18.25 | 3 | <0.001 | 392.99 |
| ASCTU | 70 | Carbon % | C~1 | NA | NA | 1 | NA | 163.17 |
| LESCA | 99 | Carbon % | C~poly(species richness, 2) | 0.08 | 4.12 | 3 | 0.019 | 150.63 |
| LUPPE | 121 | Carbon % | C~1 | NA | NA | 1 | NA | 261.78 |
| PANVI | 49 | Carbon % | C~1 | NA | NA | 1 | NA | 113.15 |
| SCHSC | 76 | Carbon % | C~1 | NA | NA | 1 | NA | 145.59 |
| SOLRI | 50 | Carbon % | C~1 | NA | NA | 1 | NA | 131.3 |
| SORNU | 27 | Carbon % | C~poly(species richness, 2) | 0.4 | 8.03 | 3 | 0.002 | 57.36 |
| ACHMI | 49 | Nitrogen % | N~poly(species richness, 2) | 0.61 | 35.24 | 3 | <0.001 | -2.71 |
| ANDGE | 162 | Nitrogen % | N~poly(species richness, 2) | 0.4 | 53.59 | 3 | <0.001 | -19.3 |
| ASCTU | 70 | Nitrogen % | N~1 | NA | NA | 1 | NA | -21.8 |
| LESCA | 99 | Nitrogen % | N~1 | NA | NA | 1 | NA | 10.39 |
| LUPPE | 121 | Nitrogen % | N~poly(species richness, 2) | 0.11 | 7.65 | 3 | <0.001 | 15.65 |
| PANVI | 49 | Nitrogen % | N~1 | NA | NA | 1 | NA | 9.9 |
| SCHSC | 76 | Nitrogen % | N~poly(species richness, 2) | 0.16 | 7.21 | 3 | 0.001 | -9.96 |
| SOLRI | 50 | Nitrogen % | N~1 | NA | NA | 1 | NA | -14.35 |
| SORNU | 27 | Nitrogen % | N~1 | NA | NA | 1 | NA | -10.36 |
| ACHMI | 49 | Non-structural carbohydrates % | NSC~1 | NA | NA | 1 | NA | 284.16 |
| ANDGE | 162 | Non-structural carbohydrates % | NSC~poly(species richness, 2) | 0.28 | 31.03 | 3 | <0.001 | 927.29 |
| ASCTU | 70 | Non-structural carbohydrates % | NSC~1 | NA | NA | 1 | NA | 411.52 |
| LESCA | 99 | Non-structural carbohydrates % | NSC~1 | NA | NA | 1 | NA | 519.77 |
| LUPPE | 121 | Non-structural carbohydrates % | NSC~1 | NA | NA | 1 | NA | 686.43 |
| PANVI | 49 | Non-structural carbohydrates % | NSC~1 | NA | NA | 1 | NA | 325.06 |
| SCHSC | 76 | Non-structural carbohydrates % | NSC~1 | NA | NA | 1 | NA | 478.6 |
| SOLRI | 50 | Non-structural carbohydrates % | NSC~1 | NA | NA | 1 | NA | 254.93 |
| SORNU | 27 | Non-structural carbohydrates % | NSC~species richness | 0.36 | 14.24 | 2 | <0.001 | 158.97 |
| ACHMI | 49 | Hemicellulose % | Hemicell~species richness | 0.1 | 5.45 | 2 | 0.024 | 253.58 |
| ANDGE | 162 | Hemicellulose % | Hemicell~poly(species richness, 2) | 0.21 | 20.54 | 3 | <0.001 | 843.33 |
| ASCTU | 70 | Hemicellulose % | Hemicell~1 | NA | NA | 1 | NA | 430.79 |
| LESCA | 99 | Hemicellulose % | Hemicell~1 | NA | NA | 1 | NA | 591.09 |
| LUPPE | 121 | Hemicellulose % | Hemicell~1 | NA | NA | 1 | NA | 700.61 |

| | | | | | | | | |
|---|---|---|---|---|---|---|---|---|
| PANVI | 49 | Hemicellulose % | Hemicell~1 | NA | NA | 1 | NA | 282.56 |
| SCHSC | 76 | Hemicellulose % | Hemicell~1 | NA | NA | 1 | NA | 459.97 |
| SOLRI | 50 | Hemicellulose % | Hemicell~1 | NA | NA | 1 | NA | 257.29 |
| SORNU | 27 | Hemicellulose % | Hemicell~species richness | 0.35 | 13.44 | 2 | 0.001 | 143.54 |
| ACHMI | 49 | Cellulose % | Cellulose~species richness | 0.23 | 14.24 | 2 | <0.001 | 267.39 |
| ANDGE | 162 | Cellulose % | Cellulose~poly(species richness, 2) | 0.39 | 51.1 | 3 | <0.001 | 817.13 |
| ASCTU | 70 | Cellulose % | Cellulose~species richness | 0.18 | 14.66 | 2 | <0.001 | 352.07 |
| LESCA | 99 | Cellulose % | Cellulose~1 | NA | NA | 1 | NA | 446.16 |
| LUPPE | 121 | Cellulose % | Cellulose~1 | NA | NA | 1 | NA | 666.25 |
| PANVI | 49 | Cellulose % | Cellulose~species richness | 0.11 | 5.52 | 2 | 0.023 | 297.75 |
| SCHSC | 76 | Cellulose % | Cellulose~1 | NA | NA | 1 | NA | 433 |
| SOLRI | 50 | Cellulose % | Cellulose~1 | NA | NA | 1 | NA | 251.73 |
| SORNU | 27 | Cellulose % | Cellulose~1 | NA | NA | 1 | NA | 135.97 |
| ACHMI | 49 | Lignin % | Lignin~1 | NA | NA | 1 | NA | 160.95 |
| ANDGE | 162 | Lignin % | Lignin~poly(species richness, 2) | 0.15 | 13.68 | 3 | <0.001 | 481.37 |
| ASCTU | 70 | Lignin % | Lignin~poly(species richness, 2) | 0.12 | 4.45 | 3 | 0.015 | 304.12 |
| LESCA | 99 | Lignin % | Lignin~1 | NA | NA | 1 | NA | 416.73 |
| LUPPE | 121 | Lignin % | Lignin~1 | NA | NA | 1 | NA | 481.21 |
| PANVI | 49 | Lignin % | Lignin~1 | NA | NA | 1 | NA | 155.7 |
| SCHSC | 76 | Lignin % | Lignin~1 | NA | NA | 1 | NA | 285 |
| SOLRI | 50 | Lignin % | Lignin~1 | NA | NA | 1 | NA | 160.07 |
| SORNU | 27 | Lignin % | Lignin~1 | NA | NA | 1 | NA | 78.42 |
| ACHMI | 49 | Chlorophyll μmol.m$^{-2}$ | Chl~poly(species richness, 2) | 0.61 | 36.21 | 3 | <0.001 | 554.24 |
| ANDGE | 162 | Chlorophyll μmol.m$^{-2}$ | Chl~species richness | 0.15 | 27.16 | 2 | <0.001 | 1906.2 |
| ASCTU | 70 | Chlorophyll μmol.m$^{-2}$ | Chl~species richness | 0.06 | 4.22 | 2 | 0.044 | 781.42 |
| LESCA | 99 | Chlorophyll μmol.m$^{-2}$ | Chl~1 | NA | NA | 1 | NA | 1096.89 |
| LUPPE | 121 | Chlorophyll μmol.m$^{-2}$ | Chl~1 | NA | NA | 1 | NA | 1393.09 |
| PANVI | 49 | Chlorophyll μmol.m$^{-2}$ | Chl~1 | NA | NA | 1 | NA | 542.03 |
| SCHSC | 76 | Chlorophyll μmol.m$^{-2}$ | Chl~1 | NA | NA | 1 | NA | 834.15 |
| SOLRI | 50 | Chlorophyll μmol.m$^{-2}$ | Chl~poly(species richness, 2) | 0.19 | 5.48 | 3 | 0.007 | 532.94 |
| SORNU | 27 | Chlorophyll μmol.m$^{-2}$ | Chl~1 | NA | NA | 1 | NA | 312.95 |
| ACHMI | 49 | β-Carotene ratio | bCar~species richness | 0.12 | 6.5 | 2 | 0.0141 | -342.08 |
| ANDGE | 162 | β-Carotene ratio | bCar~species richness | 0.06 | 9.75 | 2 | 0.0021 | -970.96 |
| ASCTU | 70 | β-Carotene ratio | bCar~species richness | 0.07 | 5.22 | 2 | 0.025 | -410.28 |
| LESCA | 99 | β-Carotene ratio | bCar~1 | NA | NA | 1 | NA | -627.79 |
| LUPPE | 121 | β-Carotene ratio | bCar~1 | NA | NA | 1 | NA | -848.13 |
| PANVI | 49 | β-Carotene ratio | bCar~1 | NA | NA | 1 | NA | -345.07 |
| SCHSC | 76 | β-Carotene ratio | bCar~species richness | 0.16 | 14.59 | 2 | <0.001 | -500.33 |
| SOLRI | 50 | β-Carotene ratio | bCar~1 | NA | NA | 1 | NA | -276.96 |
| SORNU | 27 | β-Carotene ratio | bCar~1 | NA | NA | 1 | NA | -159.93 |
| ACHMI | 49 | Lutein ratio | Lut~poly(species richness, 2) | 0.3 | 9.79 | 3 | <0.001 | -326.52 |

| | | | | | | | | |
|---|---|---|---|---|---|---|---|---|
| ANDGE | 162 | Lutein ratio | Lut~species richness | 0.14 | 25.03 | 2 | <0.001 | 1216.69 |
| ASCTU | 70 | Lutein ratio | Lut~1 | NA | NA | 1 | NA | -451.44 |
| LESCA | 99 | Lutein ratio | Lut~1 | NA | NA | 1 | NA | -708.36 |
| LUPPE | 121 | Lutein ratio | Lut~1 | NA | NA | 1 | NA | -845.29 |
| PANVI | 49 | Lutein ratio | Lut~poly(species richness, 2) | 0.14 | 3.86 | 3 | 0.0283 | -389.26 |
| SCHSC | 76 | Lutein ratio | Lut~poly(species richness, 2) | 0.2 | 9.21 | 3 | <0.001 | -524.97 |
| SOLRI | 50 | Lutein ratio | Lut~1 | NA | NA | 1 | NA | -288.15 |
| SORNU | 27 | Lutein ratio | Lut~1 | NA | NA | 1 | NA | -217.1 |
| ACHMI | 49 | Xanthophyll pool ratio | VAZ~1 | NA | NA | 1 | NA | -304.94 |
| ANDGE | 162 | Xanthophyll pool ratio | VAZ~1 | NA | NA | 1 | NA | -845.1 |
| ASCTU | 70 | Xanthophyll pool ratio | VAZ~1 | NA | NA | 1 | NA | -343.1 |
| LESCA | 99 | Xanthophyll pool ratio | VAZ~poly(species richness, 2) | 0.14 | 8.04 | 3 | <0.001 | -595.48 |
| LUPPE | 121 | Xanthophyll pool ratio | VAZ~1 | NA | NA | 1 | NA | -684.25 |
| PANVI | 49 | Xanthophyll pool ratio | VAZ~1 | NA | NA | 1 | NA | -285.09 |
| SCHSC | 76 | Xanthophyll pool ratio | VAZ~poly(species richness, 2) | 0.16 | 7.1 | 3 | 0.0015 | -440.86 |
| SOLRI | 50 | Xanthophyll pool ratio | VAZ~poly(species richness, 2) | 0.17 | 4.85 | 3 | 0.012 | -257.05 |
| SORNU | 27 | Xanthophyll pool ratio | VAZ~1 | NA | NA | 1 | NA | -143.09 |
