## appendix S1 for "Coupling spectral and resource-use complementarity in experimental grassland and forest communities"

### **Appendix S1**. Do species' foliar traits change depending on the diversity level of the plant community?

The degree to which species occupy unique hypervolumes in trait space depends on intraspecific trait variation, which, in turn, influences species' spectral variation. In the BioDIV experiment, located in a flat and fenced area with homogenous sandy soil, we expect that the diversity level (species richness) of the plant community influences species' traits because a focal individual's neighbours modify the light environment, microclimate and water access, important factors driving intra-specific trait variation. We tested six alternative hypotheses concerning the effect of the diversity of the plant community on the variability of species' foliar traits and selected the most plausible model based on Akaike's information criterion (AIC): (1) traits do not differ among species and do not change with increasing species richness (formula: trait ~ 1); (2) traits change with increasing species richness but do not differ among species (formula: trait ~ species richness); (3) traits differ among species but do not change with increasing species richness (formula: trait ~ species), such that y-intercepts can vary but the slope is always zero; (4) traits differ among species but changes with increasing species richness are consistent among species (formula: trait ~ species + species richness), such that the y-intercepts can vary but the slope is the same; (5) traits do not differ among species in monoculture but can vary among species with increasing species richness (formula: trait ~ species richness/species), such that the y-intercept is the same but the slopes can vary; (6) traits can vary among species and change for each species differently with increasing species richness (formula: trait ~ species richness*species), such that y-intercepts and slopes can vary.

We sampled spectral and foliar trait data in 35 plots covering the diversity levels of one, two, four, eight and 16 species per plot (for number of individuals per species see Material and Methods). Here we only included species that were found in at least four out of the five diversity levels of the experiment. We found that for all foliar traits, except lignin content, the best fitting models included an interaction effect between species identity and species richness (see electronic supplementary material, table S6). This means that foliar traits varied among species and that trait changes with the diversity level of the plant community were species-specific. We thus modelled trait changes for each species separately and found support for traits remaining relatively stable across diversity levels (species-specific intercept only models, trait~1) and traits changing with the diversity level of the plant community, with support for both linear and quadratic relationships depending on the trait and species considered (see electronic supplementary material, table S7 and figure S12).
